# STING-dependent cytosolic DNA sensing pathway drives the progression to leukemia in TET2-mutated HSPCs

**DOI:** 10.1101/2022.12.17.520899

**Authors:** Jiaying Xie, Mengyao Sheng, Shaoqin Rong, Chao Wang, Wanling Wu, Jingru Huang, Yue Sun, Pingyue Chen, Yushuang Wu, Yuanxian Wang, Lan Wang, Bo O. Zhou, Xinxin Huang, Colum P. Walsh, Stefan K. Bohlander, Jian Huang, Xiaoqin Wang, Hai Gao, Dan Zhou, Yuheng Shi, Guo-Liang Xu

## Abstract

Somatic loss-of-function mutations of the dioxygenase Ten-eleven translocation-2 (TET2) occur frequently in individuals with clonal hematopoiesis (CH) and acute myeloid leukemia (AML). These common hematopoietic disorders can be recapitulated in mouse models. However, the underlying mechanisms by which the deficiency in TET2 promotes these disorders remain largely unknown. Here we show that the cyclic guanosine monophosphate-adenosine monophosphate synthase (cGAS)-stimulator of interferon genes (STING) pathway is activated to mediate the effect of TET2 deficiency in leukemogenesis in mouse models. DNA damage arising in *Tet2*-deficient hematopoietic stem/progenitor cells (HSPCs) leads to activation of the cGAS-STING pathway which in turn induces the development of CH and myeloid transformation. Notably, both pharmacological inhibition and genetic deletion of STING suppresses *Tet2* mutation-induced aberrant myelopoiesis. In patient-derived xenograft (PDX) models, STING inhibition specifically attenuates the proliferation of leukemia cells from TET2-mutated individuals. These observations suggest that the hematopoietic transformation associated with TET2 mutations is powered through sterile inflammation dependent on the activated cGAS-STING pathway, and that STING may represent a potential target for intervention of relevant hematopoietic malignancies.

**Key points:**

- *Tet2* deficiency leads to DNA damage which in turn activates the cGAS-STING pathway to induce an inflammatory response
- Blocking STING in TET2-mutated hematopoietic stem/progenitor cells suppresses clonal hematopoiesis in mice and leukemogenesis in patient-derived xenograft models

## Introduction

TET2 is a dioxygenase that catalyzes the three steps of oxidation of methylated cytosines to facilitate demethylation of genomic DNA.^1^ As an epigenetic regulator, TET2 plays important roles in various physiological and pathological processes involving cell fate determination and cancer development. It is essential for the homeostasis of hematopoietic stem/progenitor cells and in the hematopoietic hierarchy and loss-of-function mutations of *TET2* are prominent drivers of age-related clonal hematopoiesis (CH) in humans.^2^ Importantly, large cohort sequencing data suggest that individuals with *TET2* mutations are also at a high risk of developing hematologic malignancies, such as myelodysplastic syndrome (MDS), myeloproliferative neoplasms (MPN) and acute myeloid leukemia (AML).^3-6^ *TET2* loss-of-heterozygosity and somatic mutations occur in nearly 40% of MDS and MPN patients, and 32% of AML patients harbor *TET2* mutations.^7^

Consistent with clinical observations, mice deficient for *Tet2* show myeloid transformation. Genetic inactivation of *Tet2* in the mouse hematopoietic system disturbed the homeostasis of HSPCs and resulted in aberrant myeloid maturation.^8-11^ In particular, *Tet2* deletion altered the HSC compartment by inducing expansion of pre-leukemic lineage^-^Sca1^+^cKit^+^(LSK) cells and changed their differentiation potential by skewing it toward monocytic/granulocytic lineages.^9^ Loss of *Tet2* in HSPCs significantly increased their replating capacity *in vitro* and self-renewal in competitive bone marrow transplantation (cBMT) assays.^10^ However, unlike aggressive AML models induced by for example *KMT2A* fusion genes,^12^ *Tet2*-deficient mice showed a long latency before developing leukemia.^9,10^ Genetic studies of AML patients indicated that mutations in *TET2* were often acquired as one of the earliest mutations.^13^ In patients with MPN or AML, *TET2* was found to co-mutate with *JAK2V617F, ASXL1, SRSF2, SF3B1, NPM1, FLT3* and *DNMT3A*.^13,14^

Recent studies demonstrated that inflammatory signals accelerate leukemogenesis driven by *TET2* loss. In *Tet2*-deficient mice, bacterial invasion due to a dysfunctional small-intestinal barrier was found to promote pre-leukemic myeloproliferation by increasing interleukin-6 (IL-6) production.^15^ *TET2* loss was shown to confer a proliferative advantage on HSPCs after exposure to TNFα and IFNγ as compared to their wildtype HSPCs.^16^ In response to inflammatory stress induced by lipopolysaccharide (LPS), *Tet2*-deficient HSPCs exhibited an increased resistance to apoptosis as well as enhanced production of IL-6 and hyperactivation of the Shp2-Stat3 signaling pathway, compared to wildtype HSPCs.^17^ Targeting SHP2 or STAT3 with inhibitors suppressed the survival advantage of *Tet2*-deficient HSPCs.^17^ These data implicate inflammation in the development of CH induced by TET2 deficiency.

Although previous animal models have faithfully recapitulated the clinical characteristics of CH and the leukemogenesis attributed to *TET2* mutations, the mechanism connecting these two remains elusive. To identify key factors responsible for this pathogenic process, we performed transcriptome analysis of distinct HSPC populations in the hematopoietic systems of *Tet2* conditional knockout (hereafter named *Tet2*^*-/-*^) mouse models. We found that the cGAS-STING pathway was involved in CH and AML development caused by mutated TET2. Deleting STING efficiently inhibited the expansion of LSK cells and the skewed myeloid differentiation in *Tet2*^*-/-*^ mice. Our findings suggest a novel mechanism that activation of cGAS-STING pathway by which TET2 loss mediates the progression of CH to leukemia, and that targeting STING might be a valid approach to delay or even prevent malignant transformation in healthy adults with CH who harbor TET2 mutations.

## Methods

### Mouse models

*Tet2*^*f/f*^ mice were generated as previously described^18^. *Sting*^-/-^ mice were kindly provided by Prof. Zhigang Lu. *Mx1-Cre* mice were purchased from Jackson Laboratory (strain # 003556). *Tet2*^*f/f*^, *Mx1-Cre* and *Sting*^*-/-*^ mice were crossed to produce *Tet2*^*f/f*^, *Mx1-Cre* and *Tet2*^*f/f*^, *Sting*^*-/-*^, *Mx1-Cre* mice. To induce *Tet2* conditional knock-out, the Cre transgene was induced in 4-week-old mice using five doses of poly(I:C) at 250 μg/body administered i.p. every other day. All mice were bred on a C57BL/6 genetic background. CD45.1 recipient mice (B6.SJL) were kindly provided by Prof. Xiaolong Liu. Immuno-deficient (B-NDG) mice were obtained from Biocytogen (Beijing, Cat. No. 110586) and used for establishing AML PDX models. All animal experiments described in this study were carried out according to the ethical guidelines of the Institute of Biochemistry and Cell Biology, Chinese Academy of Sciences, China.

### Bone marrow transplantation assays

Total BM cells were isolated respectively from *Tet2*^*f/f*^; *Sting*^*-/-*^; *Tet2*^*f/f*^, *Mx1-Cre*; *Tet2*^*f/f*^, *Sting*^*-/-*^, *Mx1-Cre* donor mice (CD45.2^+^). For non-competitive BM transplantations, 1 × 10^6^ CD45.2^+^ donor BM cells were transplanted into lethally irradiated (9.5Gy) 8-10 weeks old CD45.1^+^ recipient mice through tail vein injection. For serial transplantations, 1 × 10^6^ BM cells were isolated from the previous recipients 18 weeks post-operatively and transplanted into lethally irradiated CD45.1^+^ recipients. For competitive transplantations, BM cells from *Tet2*^*f/f*^; *Sting*^*-/-*^; *Tet2*^*f/f*^, *Mx1-Cre*; *Tet2*^*f/f*^, *Sting*^*-/-*^, *Mx1-Cre* donor mice (CD45.2^+^) and competitor mice (CD45.1^+^; B6.SJL) were mixed at a 1:1 ratio (0.5×10^6^ cells each) and injected into the tail vein of lethally irradiated (9.5Gy) CD45.1^+^ recipient mice. Tail vein blood was collected from the recipient mice every four weeks after transplantation, and the nucleated blood cells were stained with antibodies against CD45.2, CD45.1, Mac1, Gr-1, CD4 and CD8 for FACS analysis.

### Human subjects

Diagnostic bone marrow was obtained from patients with AML treated at the Huashan Hospital of Fudan University. All cells were collected after obtaining informed consent from AML patients and the study followed the ethical guidelines of Huashan Hospital of Fudan University. Human CD34^+^ cells were isolated from cord blood of healthy donors after obtaining informed consent.

### Patient-derived xenografts

3 × 10^6^ mononuclear cells (MNCs) from AML patients were transplanted into sub-lethally irradiated (1.8Gy) B-NDG mice. When the average engraftment of hCD45 cells in PB reached 1%, mice were treated with either DMSO (corn oil) or 2 μmol/body H-151 every other day for 30 days. %hCD45 engraftment in PB was analyzed after 5 doses, and mice were sacrificed at day 30 to assess the leukemia burden in BM.

### Additional methods

Additional methods including lentivirus preparation, colony formation assay, transduction of stem cells, flow cytometric analysis, immunofluorescence assay, PCR and real-time qPCR analysis, RNA library preparation and analysis, extraction and quantification of cGAMP, comet assay and statistical methods are provided in supplemental methods.

## Results

### Activation of STING pathway in *Tet2-*deficient HSPCs

To avoid the interference from acute inflammatory stress potentially caused by poly(I:C) administration that was used for the induction of *Mx1*-Cre mediated gene knockout, mice were maintained for 8 weeks post induction before being sacrificed for analysis. We performed RNA-seq of hematopoietic stem cells and lineage-restricted progenitor cells in bone marrow mononuclear cells (BM-MNCs) of wildtype (WT) and conditional knockout mice (supplemental Figure 1A-B). The pathway enrichment analysis of DEGs revealed the involvement of type I interferon response specifically enriched in *Tet2*^*-/-*^ long term hematopoietic stem cells (LT-HSCs), short term hematopoietic stem cells (ST-HSCs) (Figure 1A) and megakaryocyte-erythroid progenitor cells (MEP) (supplemental Figure 1C), suggesting that the innate immunity-related pathways are activated in these cells. Among the upregulated innate immunity-related genes, STING,^19-21^ the key adaptor of the cytosolic DNA sensing pathway (cGAS-STING pathway),^22^ showed an expression change greater than 2-fold in *Tet2*^*-/-*^ HSCs compared to WT cells, especially in LT-HSCs (Figures 1B and supplemental Figure 1D-F). Given that the cGAS-STING pathway is a key component of the innate immune system that detects cytosolic double-stranded DNA (dsDNA) and stimulates inflammatory genes expression,^23^ we reasoned that the upregulation of innate immunity-related genes in *Tet2*^*-/-*^ HSCs might be due to the activation of that pathway.

**Figure 1.**
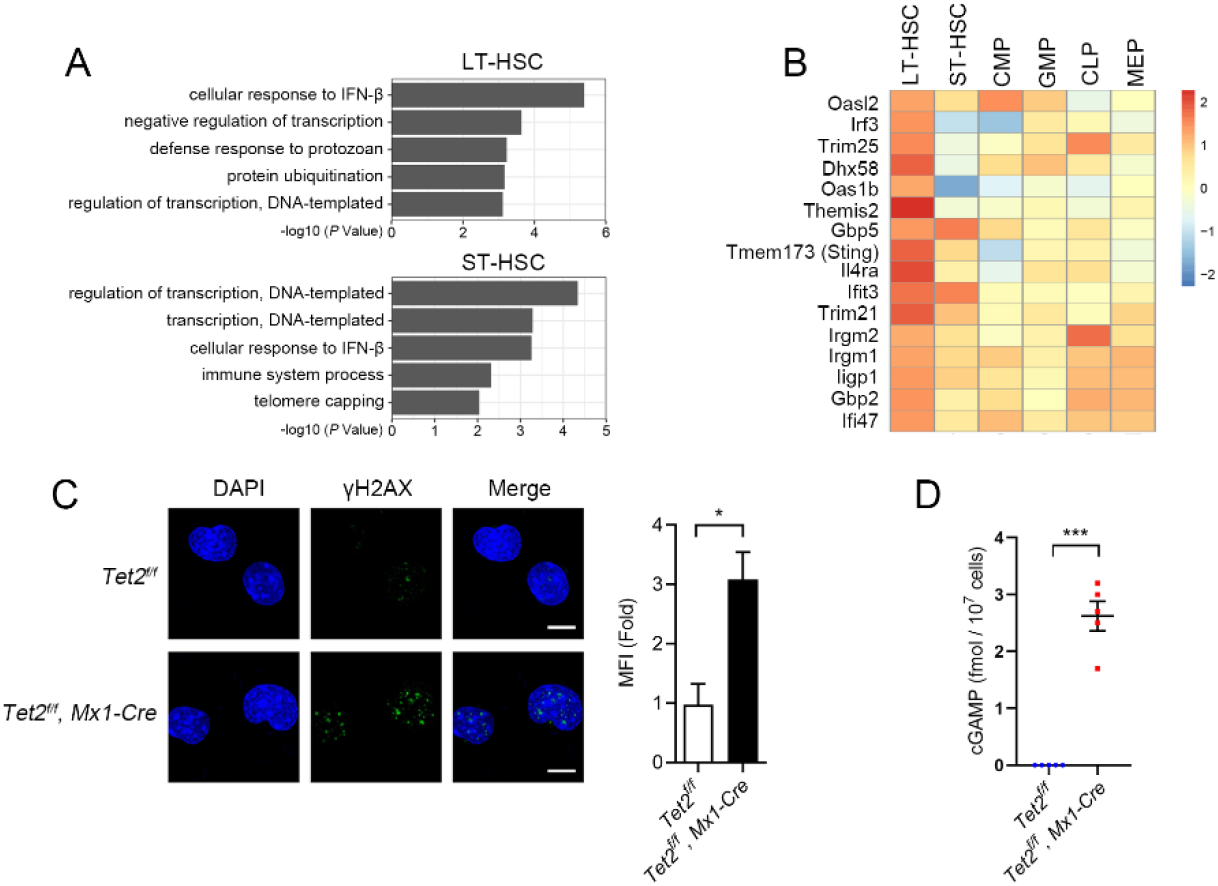
cGAS-STING pathway is activated in *Tet2*^*-/-*^ mouse HSPCs. (A) Gene sets enrichment analysis of differentially expressed genes showing activation of type I interferon response in *Tet2*^*-/-*^ hematopoietic stem cells (HSCs) compared to WT HSCs (*n* = 4 mice, *p* < 0.05). (B) Heatmap showing activation of interferon- and inflammation-related genes in *Tet2*^*-/-*^ LT-HSCs (*P* < 0.05). Scale bar denotes log2-transformed fold change. Four replicates were analyzed for each cell type. (C) Immunofluorescence of *Tet2*^*-/-*^ LSK cells stained with γH2AX antibody (green). The quantification of γH2AX signal is shown to the right (MFI: Mean fluorescence intensity; scale bar, 5μm). (D) cGAMP level in *Tet2*^*-/-*^ BM-MNCs quantified by LC-MS (*n* = 5 mice). LT-HSC, long term hematopoietic stem cell; ST-HSC, short term HSC; CMP, common myeloid progenitor; GMP, granulocyte macrophage progenitor; CLP, common lymphoid progenitor; MEP, megakaryocyte erythroid progenitor. Statistical significance was assessed by t-test (C, D). Data are mean ± s.e.m., \**P* < 0.05; \*\*\**P* < 0.005.

While infections are usually absent in the bone marrow (BM) environment of specific-pathogen-free mice, cellular DNA arising from genome instability represents a class of damage-associated molecular patterns (DAMP) which could activate the cGAS-STING pathway and stimulate the type I interferon response.^24-26^ Previous studies have shown that the deficiency of TET enzymes impairs DNA repair and induces hypermutagenicity in HSPCs.^27,28^ Indeed, double-strand breaks (DSBs) were increased in *Tet2*^*-/-*^ LSK cells compared with WT cells, as shown by anti-γH2AX immunofluorescence (Figure 1C) and comet assay (supplemental Figure 1G). This is consistent with the disturbed expression of DNA damage response- and repair-related genes in *Tet2*^*-/-*^ LT-HSCs (supplemental Figure 1H). The protein levels of STING and phosphorylated STING^29^ and TBK1^30^, two biomarkers for activation of the STING-mediated type I interferon response, were increased in *Tet2*^*-/-*^ BM-MNCs (supplemental Figure 2A-B). Moreover, the expression of *Ifnβ* and *Il-6* was upregulated in *Tet2*^*-/-*^ LSK cells (supplemental Figure 2C). To determine whether the innate immune response in *Tet2*^*-/-*^ HSPCs is dependent on STING, we knocked down *Sting* and *Mavs*,^31-34^ crucial components of their respective cytosolic dsDNA/dsRNA sensing pathways, in *Tet2*^*-/-*^ LSK cells with shRNA. Notably, only *Sting* ablation dampened the upregulation of *Ifnβ* induced by *Tet2* deficiency (supplemental Figure 2D). It is known that cGAS triggers innate immune response through synthesis of cyclic GMP-AMP (cGAMP), which binds to and activates the adaptor protein STING.^35^ We then quantified the absolute amounts of cGAMP in the BM-MNCs from WT and *Tet2*^*-/-*^ mice using liquid chromatography-mass spectrometry (LC-MS). The amount of cGAMP in *Tet2*^*-/-*^ BM-MNCs reached 3 fmol/10^7^ cells, whereas it was undetectable in WT BM-MNCs (Figures 1D and supplemental Figure 2E), confirming the activation of cGAS-STING pathway. Together, these data demonstrate that DNA damage induced by *Tet2* deficiency activates the cGAS-STING pathway, stimulating the downstream innate immune response in HSPCs.

### STING mediates skewed myelopoiesis and enhanced self-renewal of HSPCs with *Tet2* deletion

Previous studies reported that *Tet2*-deficient HSCs have a growth advantage, which can be increased by inflammatory cytokines, such as IL-6^15^ or TNF-α^36^. However, few studies to date have characterized the endogenous drivers of inflammation that contribute to the clonal expansion of *Tet2*-deficient HSPCs in the absence of an external inflammatory challenge such as microbial infection or LPS treatment. As cell-autonomous activation of the cGAS-STING pathway has been correlated with the expression of type I interferon and pro-inflammatory cytokines in *Tet2*^*-/-*^ HSPCs, it is tempting to speculate that blocking STING activation would suppress the increased self-renewal of these cells. We tested this hypothesis with the STING inhibitor C-178^37^ in a serial colony-forming unit (CFU) assay. Indeed, C-178 significantly reduced the repopulation capacity of *Tet2*^*-/-*^ LSK cells, and this effect was abrogated by IFNβ treatment (supplemental Figure 3A). The inhibitory effect of C-178 appeared specific to the LSK cells from *Tet2*^*-/-*^ mice as LSK cells from MLL-AF9^38^ or Nup98-Hox13 (NHD13)^39^ transgenic mice could not be inhibited (supplemental Figure 3B). The C-178 inhibitory effect was correlated with cGAS-STING activation in response to DNA damage, as increased γH2AX signal and elevated expression of *Ifnβ* and *Il-6* were observed in cKit^+^ *Tet2*^*-/-*^ cells but not in WT, MLL-AF9- or NHD13-harboring cells (supplemental Figure 3C-D).

Because the serial replating capacity of *Tet2*^*-/-*^ HSPCs was significantly reduced by C-178, we reasoned that blocking the cGAS-STING pathway may alleviate *Tet2* deficiency-induced expansion of HSPCs and aberrant myelopoiesis. To examine this possibility, we crossed *Tet2*^*f/f*^, *Mx1-Cre* with *Sting*^*-/-*^ to generate *Tet2*^*f/f*^, *Sting*^*-/-*^, *Mx1-Cre* mice. By administering poly(I:C) to these mice every other day for 10 days, we obtained double knockout mice for *Tet2* and *Sting* (hereafter named *Tet2*;*Sting*^*DKO*^) (Figure 2A). Loss of STING was confirmed in the *Sting*^*-/-*^ and *Tet2*;*Sting*^*DKO*^ BM-MNCs (supplemental Figure 4A). *Ifnβ* and *Il-6* were significantly downregulated in the primary cKit^+^ cells isolated from *Tet2*;*Sting*^*DKO*^ BM compared with *Tet2*^*-/-*^ cells, indicating that the cGAS-STING pathway is required for the induction of these cytokines (supplemental Figure 4B). Moreover, the growth advantage of *Tet2*^*-/-*^ cKit^+^ cells was reduced after *Sting* deletion (supplemental Figure 4C). The repopulation capacity of *Tet2*;*Sting*^*DKO*^ LSK cells was also significantly reduced compared with that of *Tet2*^*-/-*^ cells, and IFNβ treatment could restore the replating capacity of *Tet2*;*Sting*^*DKO*^ LSK cells (supplemental Figure 4D). Collectively, these data suggest that an activated cGAS-STING pathway functions to establish a local inflammatory environment which supports and stimulates the self-renewal of *Tet2*^*-/-*^ HSPCs.

**Figure 2.**
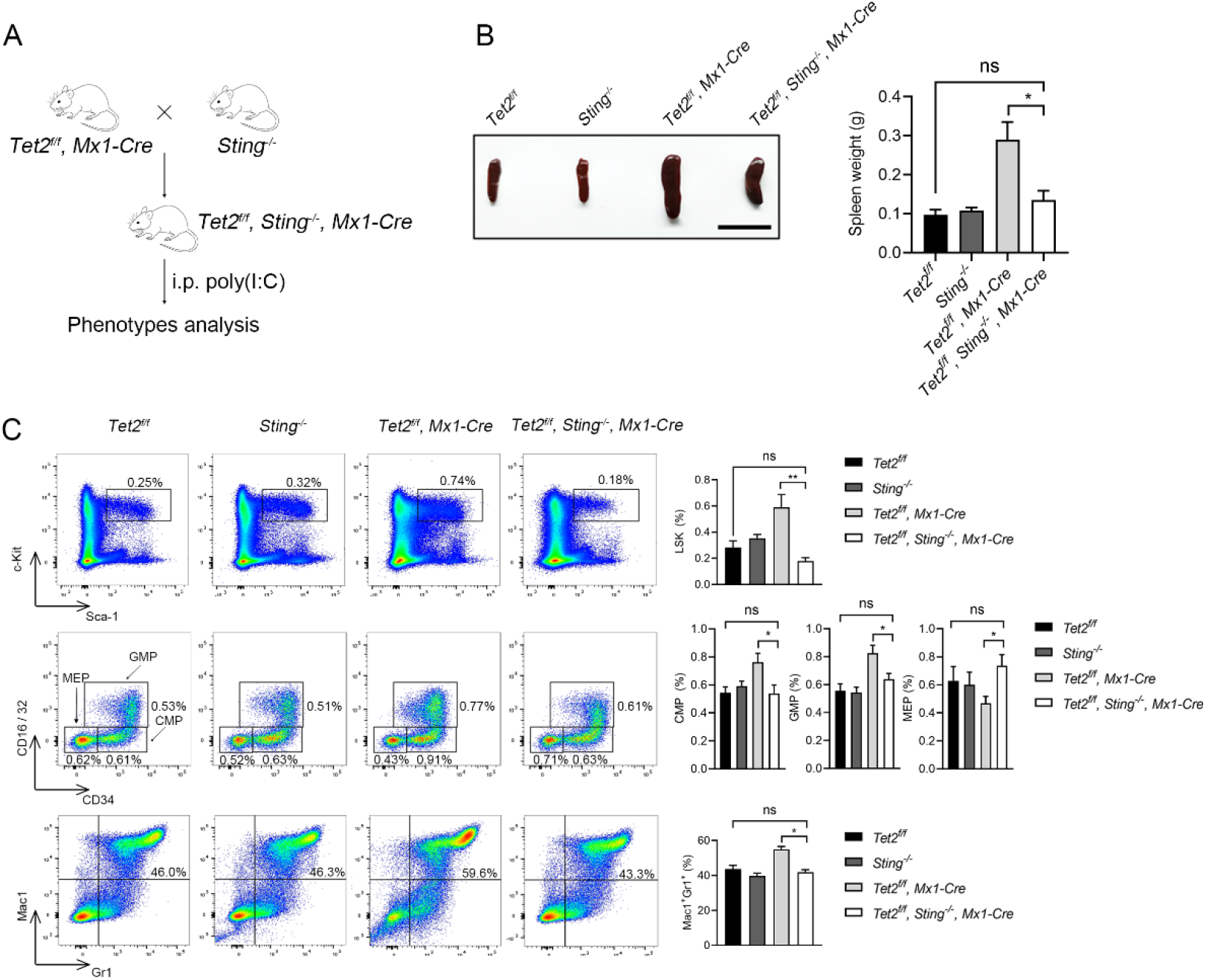
*Sting* is required for the increased self-renewal and myeloid-skewed differentiation of *Tet2*^*-/-*^ mouse HSPCs. (A) Breeding strategy for inducible deletion of *Tet2* in the hematopoietic system on the *Sting*^-/-^ background. i.p., intraperitoneal injection. (B) Comparison of spleens from representative mice of indicated genotypes. Shown on the right is the quantification of spleen weight (*n* = 5 mice; scale bar, 1cm). (C) *Sting* deletion attenuates the aberrant hematopoiesis caused by *Tet2* deficiency. The percentages of LSK cells (top panel), oligopotent progenitor cells (CMP, GMP and MEP, middle panel) and Mac1^+^Gr1^+^ myeloid cells (bottom panel) in the BM of indicated mice were determined by flow cytometric analysis. All mice were sacrificed for analysis 12 months after poly(I:C) injection. The quantification is shown to the right (*n* = 5 mice). Statistical significance was assessed by t-test (B, C). Data are mean ± s.e.m., \**P* < 0.05; \*\**P* < 0.01; “ns”: not significant.

Previous studies have shown that inactivation of *Tet2* leads to an increased LSK pool and differentiation skewing towards monocytic/granulocytic lineages.^9-11^ We then asked whether these phenotypes can be alleviated by additional *Sting* deletion. At 12 months post poly(I:C) induction, the size and weight of spleens in *Tet2*;*Sting*^*DKO*^ mice were significantly reduced compared with those from *Tet2*^*-/-*^ mice (Figure 2B). We further examined myeloid differentiation and erythropoiesis in the peripheral blood (PB). Notably, myeloid-skewing induced by *Tet2* deficiency was attenuated by *Sting* deletion (supplemental Figure 4E), and reduced erythropoiesis, another typical phenotype of *Tet2*^*-/-*^ mice,^40^ was fully restored by *Sting* deletion (supplemental Figure 4F). *Sting* loss appeared to affect specifically the *Tet2*^*-/-*^ phenotypes because *Sting*-KO alone had no discernible effect on lineage differentiation.

We observed significantly more LSK cells in the BM of *Tet2*^*-/-*^ mice. However, there was almost no change in the LSK population when both *Tet2* and *Sting* were deleted. Furthermore, targeted deletion of *Sting* rescued the alteration of oligopotent progenitor cells in *Tet2*^*-/-*^ mice, including the increase of CMP and GMP and the decrease of MEP (Figure 2C). Moreover, *Tet2*;*Sting*^*DKO*^ mice exhibited a comparable proportion of myeloid cells to *Tet2*^*f/f*^ control mice, and the expansion of myeloid cells in the BM was only observed in *Tet2*^*-/-*^ mice (Figure 2C). Thus, cGAS-STING activation is required for the *Tet2* deficiency-induced expansion of myeloid progenitor cells and *Sting* deletion reduces the skewed differentiation of oligopotent progenitor cells. Interestingly, the HSC pool was enlarged as a result of *Sting, Tet2* double deletion as evidenced by the significantly increased proportion of both ST- and LT-HSCs in the BM of *Tet2*;*Sting*^*DKO*^ mice compared with the WT mice (supplemental Figure 4G). Together, these data suggest that the defects in hematopoiesis and myeloid skewing induced by *Tet2* deficiency are mediated by STING.

### *Sting* is indispensable for *Tet2* loss induced CH and leukemogenesis

In a non-competitive transplantation assay (Figure 3A), *Tet2*^*-/-*^ BM donor cells exhibited skewed differentiation towards the myeloid lineage in the peripheral blood (PB) of recipient mice, whereas the myeloid skewing phenotype was ameliorated after *Sting* deletion (Figure 3B). Eighteen weeks after transplantation, recipients were sacrificed for analysis. Consistent with significant myeloid expansion, the populations of oligopotent progenitors (CMP, GMP, and MEP) and LSK cells from *Tet2*^*-/-*^ donors had increased more than 2-fold relative to those from *Tet2*^*f/f*^ control donors. In contrast, these cell proportions were not significantly changed in mice who had received cells *Tet2*;*Sting*^*DKO*^ donors (Figure 3C). In the recipients of cells from *Tet2*^*-/-*^ donors, the HSC proportion (ST- and LT-combined) was reduced by 60%; in contrast, recipients of cells from *Tet2*;*Sting*^*DKO*^ donors had normal-sized HSC populations (Figure 3D). We observed that the phenotype of enlarged spleen and compromised differentiation potential towards erythroid lineage seen in recipient of cells from *Tet2*^*-/-*^ donors was rescued in recipients of cells from *Tet2*;*Sting*^*DKO*^ donors (Figure 3E-F), suggesting that the *Tet2;Sting*^*DKO*^ donor cells sustained grossly normal hematopoiesis. Strikingly, *Sting* deletion significantly prolonged the survival of recipients of *Tet2*^*-/-*^ BM cells (Figure 3G), indicating that targeting STING might represent a novel therapeutic approach to inhibit the onset of leukemogenesis driven by mutated *TET2*. Taken together, the above data suggest that additional deletion of *Sting* rescues both the function and homeostasis of the hematopoietic system of *Tet2*^*-/-*^ mice.

**Figure 3.**
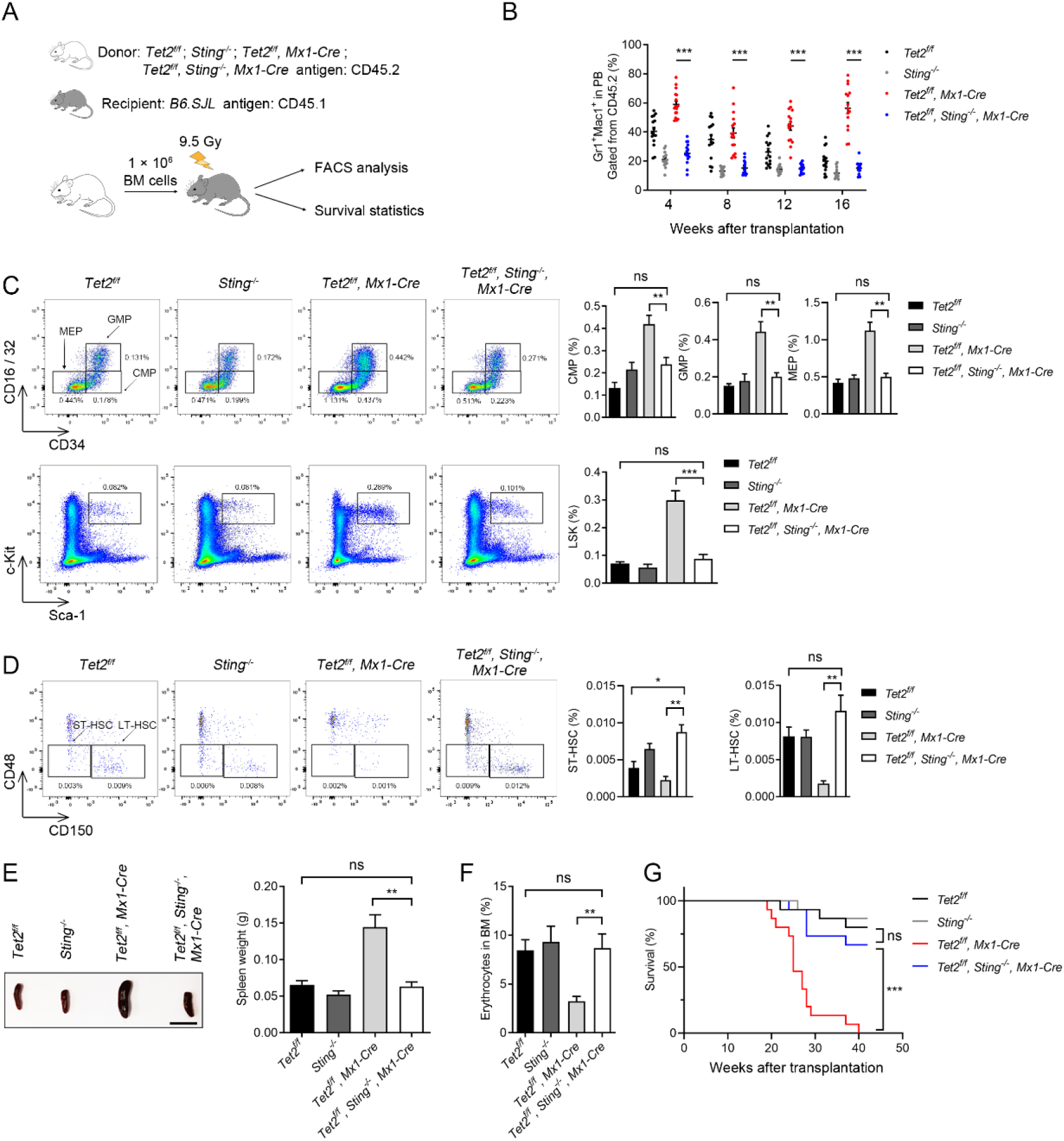
*Sting* deletion abrogates *Tet2* deficiency-induced skewed hematopoiesis in bone marrow transplantation models. (A) Schematic of bone marrow transplantation assay. (B) Myeloid skewing of *Tet2*^*-/-*^ cells after transplant (red dots) is corrected by *Sting* (blue dots) deletion. The scatter plot shows the quantification of donor-derived mature myeloid cells in the peripheral blood (PB) of primary recipients transplanted with indicated donor BM cells. (C) *Sting* deletion attenuates the expansion of *Tet2*^*-/-*^ progenitor cells in recipient mice. Representative FACS plots of progenitor cells from the BM cells of indicated donor groups are shown on the left. Top row, CMP, GMP, MEP; Bottom row, LSK. Quantification is shown on the right. (D) *Sting* deletion increases the number of *Tet2*^*-/-*^ HSCs. Representative FACS plots of HSCs from donors of indicated genotypes are shown on the left. Quantification is shown on the right. (E) Comparison of spleens from recipient mice of indicated genotypes. Shown to the right is the quantification of spleen weight. (n = 3 donors and 15 recipients for each genotype; scale bar, 1cm) (F) *Sting* deletion restores normal erythropoiesis of *Tet2*^*-/-*^ donors. The proportion of CD71^+^Ter119^+^ erythrocytes in the BM were analyzed by FACS. (G) Survival analysis of primary transplant recipients of indicated donor groups (n = 3 donors and 15 recipients for each genotype). In BM transplantation assays, all mice (n = 3 donors and 15 recipients for each genotype) were sacrificed and analyzed at 18 weeks after transplantation. Statistical significance was assessed with two-way ANOVA (B-D), t-test (E, F), and Mantel–Cox log-rank test (G). Data are mean ± s.e.m., \**P* < 0.05; \*\**P* < 0.01; \*\*\**P* < 0.005; “ns”: not significant.

To further investigate whether STING is indispensable for the increased self-renewal of *Tet2*^*-/-*^ HSPCs, serial-transplantation assays were performed (Figure 4A). A previous study showed that *Tet2*-deficient cells exhibited enhanced engrafting potential in such an assay.^41^ We performed three rounds of serial transplantation, each lasting for 16 weeks. The increase in the population of *Tet2*^*-/-*^ progenitor cells derived from donor mice was abrogated by *Sting* deletion (Figures 4B and 4C). Additionally, while *Tet2*^*-/-*^ donors generated 0.5-3.5-fold more LSK cells than *Tet2*^*f/f*^ donors in each round of transplantation, LSK cells were not increased in the recipients of cells from *Tet2*;*Sting*^*DKO*^ donors (Figure 4D). Interestingly, although the frequency of LT-HSCs usually decreased after each round of transplantation, the LT-HSC population in the recipients of *Tet2*;*Sting*^*DKO*^ donors was larger than that of the other genotypes (Figure 4E). Together, these transplantation experiments demonstrate that blocking *Sting* in *Tet2*-deficient hematopoietic cells restores normal hematopoiesis.

**Figure 4.**
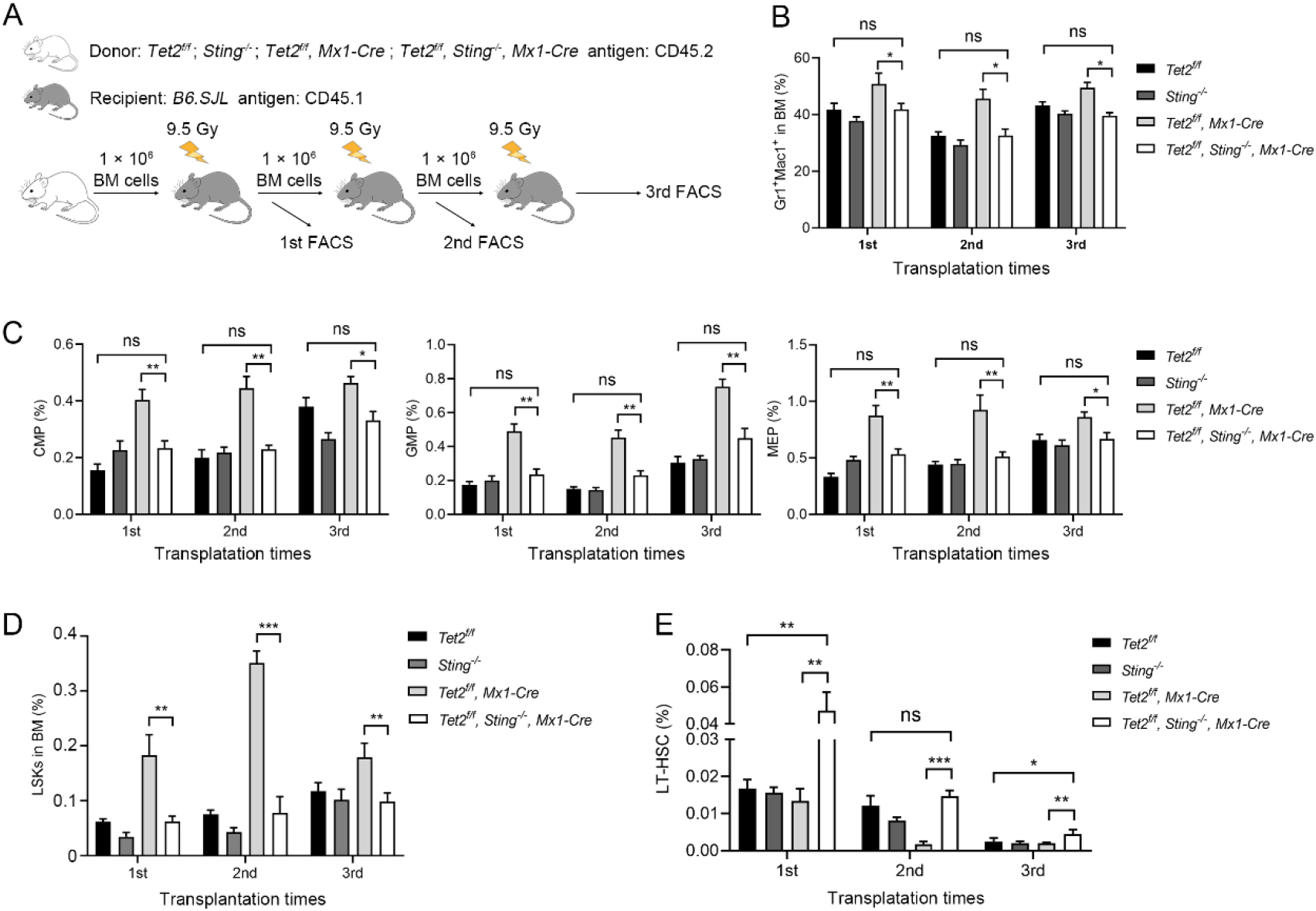
Blocking STING restores normal hematopoiesis of *Tet2*^*-/-*^ BM cells in serial transplantation models. (A) Schematic of serial transplantation assays. (B) Myeloid skewing of *Tet2*^*-/-*^ BM donors is corrected by *Sting* deletion. (C) *Sting* deletion attenuates the expansion of transplanted *Tet2*^*-/-*^ progenitor cells in recipient mice. (D) *Sting* deletion mitigates the increased expansion of *Tet2*^*-/-*^ LSK cells in serial transplantation recipients. (E) *Sting* deletion increases the proportion of *Tet2*^*-/-*^ donor LT-HSCs in serial transplantation recipients. All mice (n = 2 donors and 10 recipients for each genotype) were sacrificed and analyzed 18 weeks after transplantation. Statistical significance was assessed with two-way ANOVA (B-D) and t-test (E). Data are mean ± s.e.m., \**P* < 0.05; \*\**P* < 0.01; \*\*\**P* < 0.005; “ns”: not significant.

To investigate whether STING is essential for the enhanced self-renewal activity of *Tet2*^*-/-*^ HSPCs, we performed competitive transplantation assays (Figure 5A). As expected, *Tet2*;*Sting*^*DKO*^ BM donor cells showed a significantly reduced level of reconstitution capacity compared to *Tet2*^*-/-*^ cells (Figure 5B), indicating that *Sting* contributes to the competitive advantage conferred by *Tet2* deficiency. We further tested whether STING inhibitor can suppress the competitive advantage of *Tet2*^*-/-*^ donor cells by co-transplanting *Tet2*^*-/-*^ and CD45.1 BM cells into recipient mice. The recipients were treated with the STING inhibitor C-176.^37^ The relative contribution of *Tet2*^*-/-*^ donor cells was reduced by ∼12% after a two-week treatment with C-176 (Figure 5C). When the BM cells of recipient mice were isolated for analysis, we observed increased erythrocytes and decreased myeloid cells in the recipients treated with C-176 compared to those treated with DMSO (Figure 5D).

**Figure 5.**
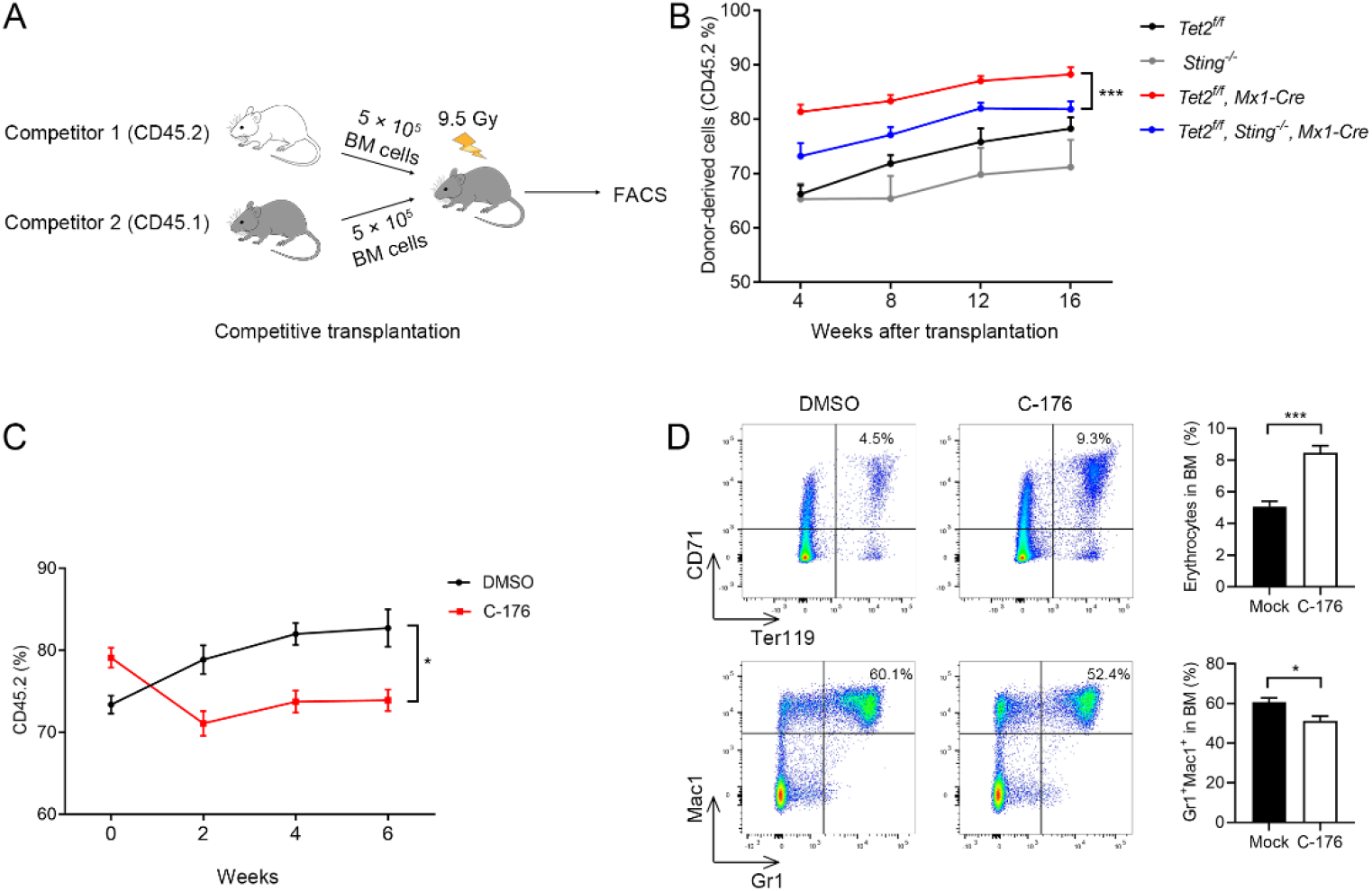
Targeting STING abrogates *Tet2* deficiency-induced clonal hematopoiesis in competitive BMT assays. (A) Schematic of cBMT assays. (B) *Sting* (blue line) deletion reduces the competitive advantage of *Tet2*^*-/-*^ donor cells (red line) in cBMT assay (n = 3 donors and 15 recipients for each genotype). (C) Sting inhibitor C-176 abrogates the competitive advantage of *Tet2*^*-/-*^ donor cells in a cBMT assay (n = 1 donor and 5 recipients). Mice were treated with C-176 or DMSO control by i.p. every 3 days and engraftment of donor cells (CD45.2) was analyzed every 2 weeks by FACS. (D) C-176 treatment promotes erythropoiesis of *Tet2*^-/-^ donor BM cells in cBMT assays (n = 1 donor and 5 recipients). Mice were treated with DMSO or C-176 by i.p. every 3 days and sacrificed after 8 weeks of inhibitor treatment. The erythropoiesis and myelopoiesis of recipients were analyzed by FACS. Statistical significance was assessed with two-way ANOVA (B-D). Data are mean ± s.e.m., \**P* < 0.05; \*\**P* < 0.01; \*\*\**P* < 0.005.

Collectively, these data demonstrate that *Sting* is indispensable for *Tet2*-loss-induced CH and leukemogenesis, and that blocking the cGAS-STING pathway by either gene deletion or small molecules can restrain skewed differentiation and malignant transformation induced by *Tet2* deficiency.

### Blocking STING suppresses AML harboring *TET2* mutation

CHs harboring *Tet2* mutation alone rarely progress to hematopoietic malignancies;^42^ cooperating mutations,^43^ such as in *Flt3* or *Jak2*^*V617F*^, accelerate leukemogenesis in mouse models.^44,45^ As the recipient mice of *Tet2*;*Sting*^*DKO*^ BM donor cells exhibited normal hematopoietic hierarchy and significantly extended survival, we wondered whether STING inhibition is able to restrain the skewed differentiation and leukemia progression of *TET2*-mutated human blood cells *in vitro* and in engrafted mice. We first examined the effect of human STING inhibitor H-151^37^ on the proliferation of human cord blood-derived CD34^+^ HSPCs upon *TET2* depletion. Consistent with a previous study^46^, ablating TET2 with shRNA increased the granulocyte-macrophage colony-forming unit (CFU-GM) but reduced the burst forming unit-erythroid (BFU-E) compared with the control group. The CD34^+^ cells with TET2 ablation showed upregulated IFNβ and increased micronuclei release (supplemental Figure 5A-B). Furthermore, the cGAS protein exactly co-localized with the micronuclei in the cytoplasm, suggesting that DNA damage induced by TET2 ablation activates the cGAS-STING pathway (supplemental Figure 5C). Notably, H-151 treatment attenuated the CFU-GM colony formation and restored erythroid colony formation of TET2-ablated cells (Figure 6A). Consistently, knockdown of STING normalized IFNβ expression and lineage-skewed colony formation in TET2-ablated CD34^+^ cells (supplemental Figure 6A-B).

**Figure 6.**
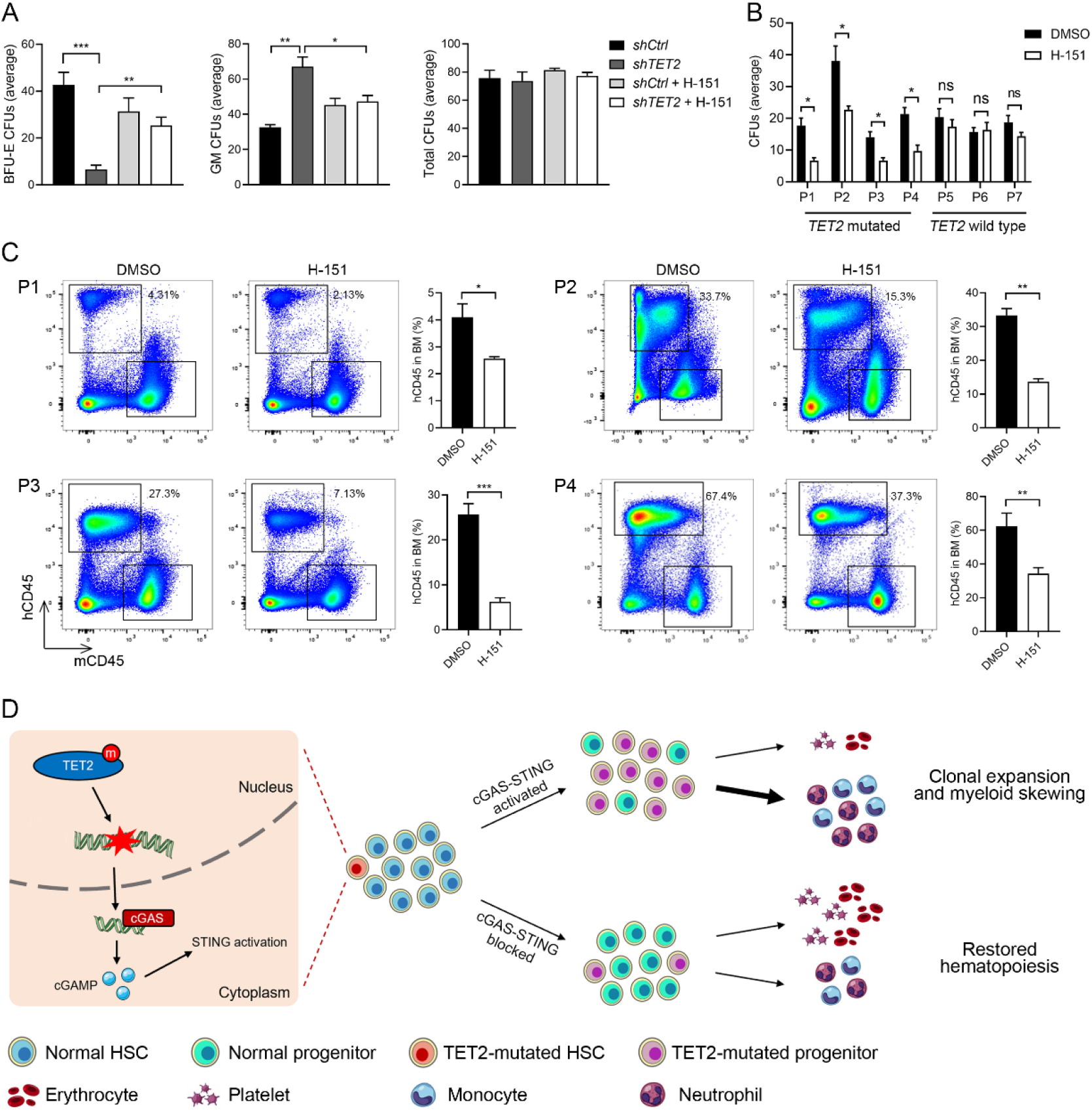
Myeloid skewing of TET2 knock-down human HSCs and leukemogenesis of TET2-mutated AML Patient cells are prevented by inhibiting STING. (A) STING inhibitor H-151 reduces myeloid colony formation and restores erythroid colony formation of TET2 knock-down cord blood CD34^+^ cells. CD34^+^ cells were infected with lentivirus carrying short hairpin RNA which targeted *TET2* (sh*TET2*) or scrambled control (sh*Ctrl*) and cultured in methylcellulose medium for 14 days. (B) H-151 inhibits colony formation of *TET2*-mutated patient AML cells. P1-4 are AML patients with *TET2* mutations, and P5-7 are those without *TET2* mutation. (C) H-151 reduces the engraftment efficiency of *TET2*-mutated patient AML cells in B-NDG mice. The percentages of human CD45-positive cells (hCD45) in total BM are indicated. (D) Cartoon summarizing our findings: HSPCs with mutated *TET2* accumulate DNA damage, which in turn activates the cGAS-STING pathway leading to the production of inflammatory cytokines. This promotes clonal expansion and myeloid differentiation skewing of the mutant HSPCs. Blocking the cGAS-STING pathway suppresses the increased self-renewal and skewed differentiation potential of the mutant HSPCs, and restores normal hematopoiesis. Statistical significance was assessed by t-test (A-C). Data are mean ± s.e.m., \**P* < 0.05; \*\**P* < 0.01; \*\*\**P* < 0.005; “ns”: not significant.

To evaluate the therapeutic potential of STING inhibition, we tested the effect of H-151 on colony formation of BM cells derived from 4 AML patients with *TET2* mutations (P1-4) and 3 AML patients without *TET2* mutations (P5-7) (supplemental Figure 7A). RT-qPCR analysis showed that *TET2* mRNA levels were reduced in P1-4 cells compared with P5-7, and the *in vitro* colony formation of P1-4 was inhibited by H-151 whereas that of P5-7 was not affected (Figure 6B and supplemental Figure 7B). We further established PDX models in immunodeficient mice using mononuclear cells from these patients. The engraftment was examined 4 weeks after transplantation, and mice were treated with H-151 from 12 weeks onwards. In the PDX models of P5-7 without *TET2* mutations, the patient-derived CD45^+^ cells continued to expand in the PB during H-151 treatment; in contrast, the proportions of P1-4-derived hCD45^+^ cells in the PB were significantly reduced by H-151 administration (supplemental Figure 7C), indicating that H-151 specifically inhibits the leukemogenesis of *TET2*-mutated patient cells. Furthermore, compared to the DMSO group, the proliferation of *TET2*-mutated leukemia cells was significantly reduced in the BM of recipient mice treated with H-151, and no inhibitory effect was observed in the recipients of leukemia cells harboring wildtype *TET2* (Figures 6C and supplemental Figure 7D). These data indicate that the leukemogenic potential of *TET2*-mutated AML mononuclear cells can be suppressed by inhibiting STING.

## Discussion

In this study, we demonstrated that DNA damage engendered by TET2 deficiency activated the cGAS-STING pathway in HSPCs. Moreover, inhibition of STING significantly attenuated the expansion of LSK and skewed myeloid differentiation in both *Tet2*^-/-^ mice and bone marrow transplant models. Compared to *Tet2*^-/-^ donor cells, *Tet2*;*Sting*^*DKO*^ donor cells restored a balanced hematopoietic hierarchy and extended significantly the lifespan of recipient mice. Leveraging AML patient-derived xenograft models, we further showed that patient-derived BM-MNCs with *TET2* mutations were more sensitive than those without *TET2* mutations to STING inhibition. Taken together, these data suggest that STING plays an essential role in hematopoietic disorders induced by *TET2* mutation and inhibition of STING represents a potential therapeutic strategy to block the production of inflammatory cytokines and mitigate the development of CH and leukemogenesis driven by *TET2* mutations (Figure 6D).

Chronic low-grade inflammation promotes myeloid differentiation and leukemogenesis in *Tet2*-deficient mice.^17,36^ Using microbial infection- or LPS injection-induced inflammatory models, several studies have shown that the hematopoietic malignancies driven by *TET2* mutation were effectively prevented by targeting the activated inflammatory pathways.^15,17,47^ These observations showed that inflammation accelerates myeloid transformation on the basis of *TET2* deficiency. However, the mechanisms by which hematopoietic disorders develop spontaneously in *Tet2* knockout specific pathogen-free mice have remained unclear. Here we show that the damaged DNA from genome instability activates the cGAS-STING pathway and induces a sterile inflammatory reaction in *Tet2*^*-/-*^ HSPCs, which in turn fosters an inflammatory environment and promotes the progression from CH to leukemia. This reveals that not only an acute inflammatory response triggered by external stimuli promotes the leukemic transformation of *TET2*-mutated HSPCs, but also that chronic sterile inflammation in mutated HSPCs can lead to an increase of the stem cell pool promoting the acquisition of additional leukemogenic mutations. It would be interesting to test whether STING is also activated in other hematopoietic malignancies involving DNA damage or genome instability, and whether targeting the cGAS-STING pathway might inhibit the malignant transformation of HSPCs.

The cGAS-STING pathway has emerged as a crucial pathway in cancer immunology. Recent studies report that activation of the cGAS-STING pathway can be used to enhance antitumor <u>immunotherapy,</u>^48,49^ in part due to the induction of type I interferons which enhance the recognition of tumor cells and the killing efficiency of T cells.^50^ However, activation of the STING pathway also enhances the IL-6-dependent survival of chromosomally unstable breast cancer cells, suggesting a pro-tumorigenic effect of cGAS-STING signaling.^51^ Therefore, the effect of an activated cGAS-STING pathway can be ambiguous. Based on our study, the stimulation of STING in *TET2*-mutated HSPCs promotes the progression to leukemia.

In summary, our study uncovers an unexpected role of STING in CH mediated by TET2 deficiency and demonstrates that targeting STING can effectively eliminate the leukemogenesis of *TET2*-mutated HSPCs. These findings provide new insights into the progression of CH to AML and an exciting new opportunity to develop therapeutic strategies to delay or stop this process.

## Supporting information

Supplement data including all supplementary figures and methods

## Acknowledgements

We acknowledge L. Wang, R. Ren, J. Chen, and Q. Wang for the discussions; J. Wang, J. Chen, and R. Su for critical reading of the manuscript; T. Chen, X. Huang and X. Wang for providing BM cells of mouse AML model, human cord blood and AML patient samples; Z. Lu for providing *Sting*^*-/-*^ mice; X. Liu for providing B6.SJL mice. SKB is supported by Leukemia & Blood Cancer New Zealand and the family of Marijana Kumerich. This work was supported by the National Natural Science Foundation of China (32000420 to D. Z.; 31901011 to Y. S.), the Shanghai Sailing Program (18YF1412200), the Natural Science Foundation of Shanghai (20ZR1408000), the Fudan University Start-up Research Grant (IDH1340038, IDH1340045, IDH1340046, IDH1340059).

## Authorship

G. X. and Y. S. conceived the original idea and, Y. S., J. X., M. S. and D. Z. designed the experiments. Experiments were performed by Y. S with the help of: M. S. for the experiments associated with FACS; J. X. for mouse modeling; J. H., Y. S. and P. C. for mouse breeding and phenotyping; W. W. for human sample collection; S. R. for the experiments associated with LC-MS; C.W. for Smart-seq2 library preparation and data analysis; Y. W. for mouse modeling and phenotyping; G. X., Y. S., D. Z., C.P.W., S.K.B., H. J. and B. Z. wrote and revised the manuscript, with contributions from all other authors.

## Supplemental Information

Supplemental information includes 6 figures and additional methods.

