## Supplement data including all supplementary figures and methods for "STING-dependent cytosolic DNA sensing pathway drives the progression to leukemia in TET2-mutated HSPCs"

Manuscripts: Jiaying Xie et al.

### Supplemental information:

Supplemental figures and additional methods are provided in supplemental data.

### Supplemental figures

Supplementary figure1

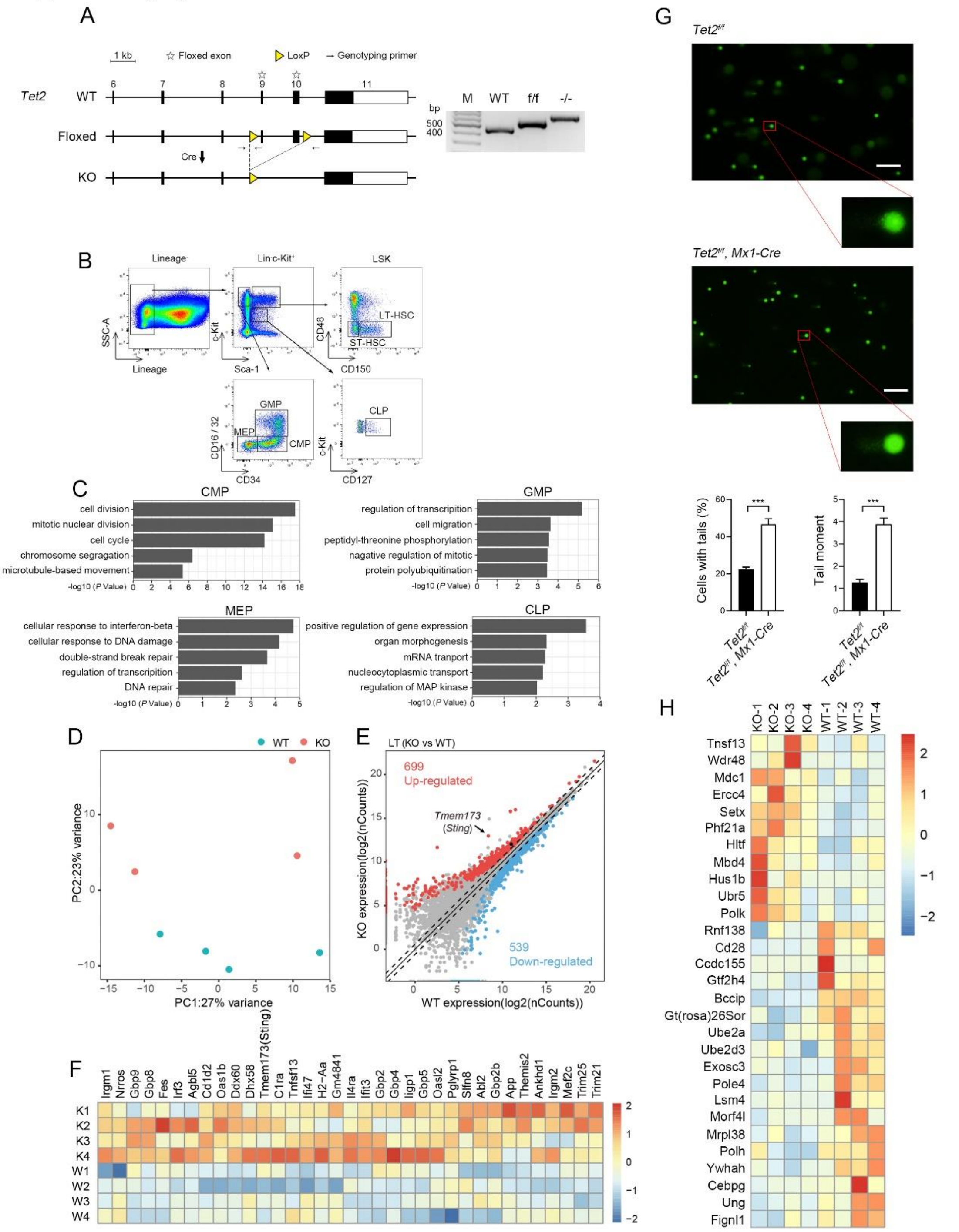


**Supplementary figure 1. Activated innate immunity pathway and increased DNA damage in hematopoietic stem cells of *Tet2*$^{-/-}$ mouse.**

(A) Schematic of Cre-mediated *Tet2* gene conditional knockout. Coding exons are shown as filled boxes. LoxP sites are shown as yellow triangles and genotyping primers as arrows. The positions of the LoxP sites flanked exons of TET2 are marked by a star. Panel to the right shows the genotyping result of Cre-mediated *Tet2* deletion.

(B) Gating strategy used for sorting of hematopoietic stem cells and oligopotent progenitor cells from bone marrows. 1) LT-HSC, $Lin^{-}cKit^{+}Sca1^{+}CD48^{-}CD150^{+}$; 2) ST-HSC, $Lin^{-}cKit^{+}Sca1^{+}CD48^{-}CD150^{-}$; 3) CMP, $Lin^{-}cKit^{+}Sca1^{-}CD34^{int/+}CD16/32^{int}$; 4) GMP, $Lin^{-}cKit^{+}Sca1^{-}CD34^{+}CD16/32^{high}$; 5) MEP, $Lin^{-}cKit^{+}Sca1^{-}CD34^{-}$ $CD16/32^{low}$; 6) CLP, $Lin^{-}cKit^{+}Sca1^{int}$ $CD127^{+}$)

(C) Enrichment analysis of differentially expressed genes (DEGs) in four groups of *Tet2*$^{-/-}$ hematopoietic progenitor cells ($n$ = 4 mice, $P < 0.05$) (see also Fig.1).

(D) PCA plot of RNA-seq data from *Tet2*$^{-/-}$ and WT LT-HSCs. Each dot represents an individual mouse ($n$ = 4 mice).

(E) Scatter plot showing the DEGs between *Tet2*$^{-/-}$ and WT LT-HSCs. Red and blue dots denote significantly changed genes ($n$ = 4 mice, $P < 0.05$), and gray dots represent unchanged genes.

(F) Heatmap showing upregulation of innate immunity- and inflammation-related genes in *Tet2*$^{-/-}$ LT-HSCs. The genes with $\log_2$(fold change) > 1 and $P < 0.05$ ($n$ = 4 mice) are shown.

(G) Comet assay showing increased DNA double strand breaks in *Tet2*$^{-/-}$ LSK cells. Quantifications of cells with tails and tail moment are shown at bottom. ($n$ = 3 mice, statistical significance was assessed by t-test, data are mean ± s.e.m., ***$P < 0.005$)

(H) Heatmap showing differential expression of DNA damage response-related genes in *Tet2*$^{-/-}$ LT-HSCs. Up- or down-regulation are defined as mean $\log_2$(fold change) > 1 and $P < 0.05$ ($n$ = 4 mice).

Supplementary figure 2

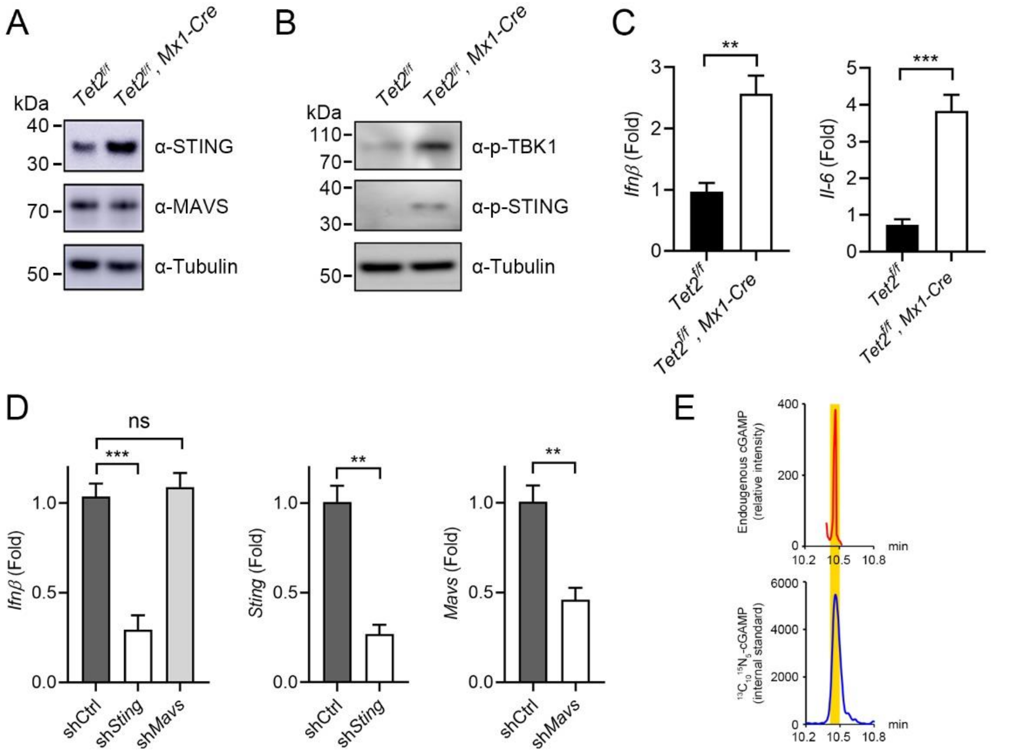


**Supplementary figure 2. The STING-mediated innate immunity pathway is activated in *Tet2*$^{-/-}$ hematopoietic cells.**

(A) Western blot showing the upregulation of *Sting* in *Tet2*$^{-/-}$ BM-MNCs. MAVS and Tubulin were used for normalization.

(B) Phosphorylation of STING and TBK1 in *Tet2*$^{-/-}$ BM-MNCs was detected by Western blot. Tubulin was used as normalization.

(C) Upregulation of *Ifnβ* and *Il-6* in *Tet2*$^{-/-}$ LSK cells. The mRNA levels of *Ifnβ* and *Il-6* are quantified by qRT-PCR (n = 3 mice)

(D) Sting is required for type I interferon production in *Tet2*$^{-/-}$ LSK cells. The mRNA levels of *Sting*, *Mavs* and *Ifnβ* were quantified by qRT-PCR (n = 3 mice).

(E) Mass chromatograms displaying quantification of intracellular cGAMP (upper panel) in *Tet2*-deficient BM-MNCs. $^{13}C_{10}{}^{15}N_5$-labeled cGAMP was spiked in the LC-MS samples as an internal standard (bottom panel) and the cGAMP peak is highlighted in yellow (n = 5 mice).

Statistical significance was assessed by t-test (C, D), data are mean ± s.e.m., ***P* <

0.01, ***$P < 0.005$, “ns”: not significant.

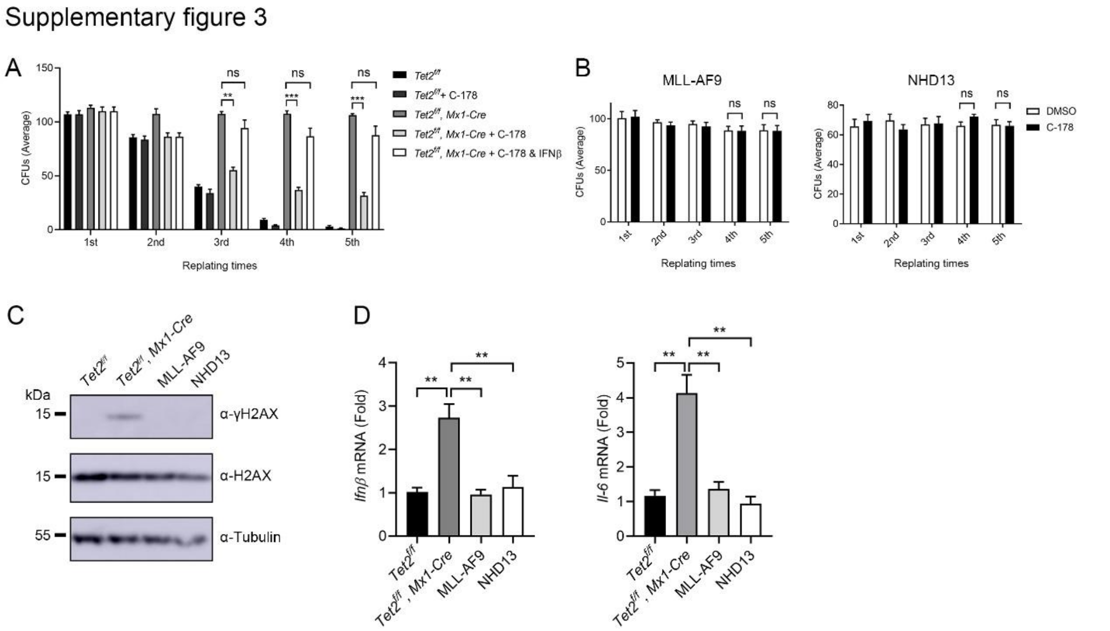


**Supplementary figure 3. STING inhibitor C-178 inhibits the replating capacity of *Tet2*$^{-/-}$ LSK cells specifically.**

(A) Reduction in the serial replating potential of *Tet2*$^{-/-}$ LSK cells cultured in the presence of C-178 (5μM). The addition of IFNβ treatment (0.2 U/ml) restored the replating ability of *Tet2*$^{-/-}$ LSK cells treated with C-178 ($n$ = 3 mice).

(B) C-178 treatment does not reduce the replating ability of MLL-AF9- or Nup98-Hox13(NHD13)-transformed LSK cells (n = 3 independent experiments). Colony formation assay was performed with LSK cells derived from AML mouse models harboring the fusion oncoproteins of MLL-AF9 (left) or Nup98-Hox13 (right).

(C) Western blot showing the phosphorylation of H2AX in *Tet2*$^{-/-}$ cKit$^{+}$ cells. H2AX and Tubulin were used as normalization

(D) Upregulation of *Ifnβ* and *Il-6* in *Tet2*$^{-/-}$ cKit$^{+}$ cells. The mRNA levels of *Ifnβ* and *Il-6* were quantified by qRT-PCR (n = 3 mice).

Statistical significance was assessed by t-test (A-E). Data are mean ± s.e.m., $^{**}P < 0.01$, $^{***}P < 0.005$, “ns”: not significant.

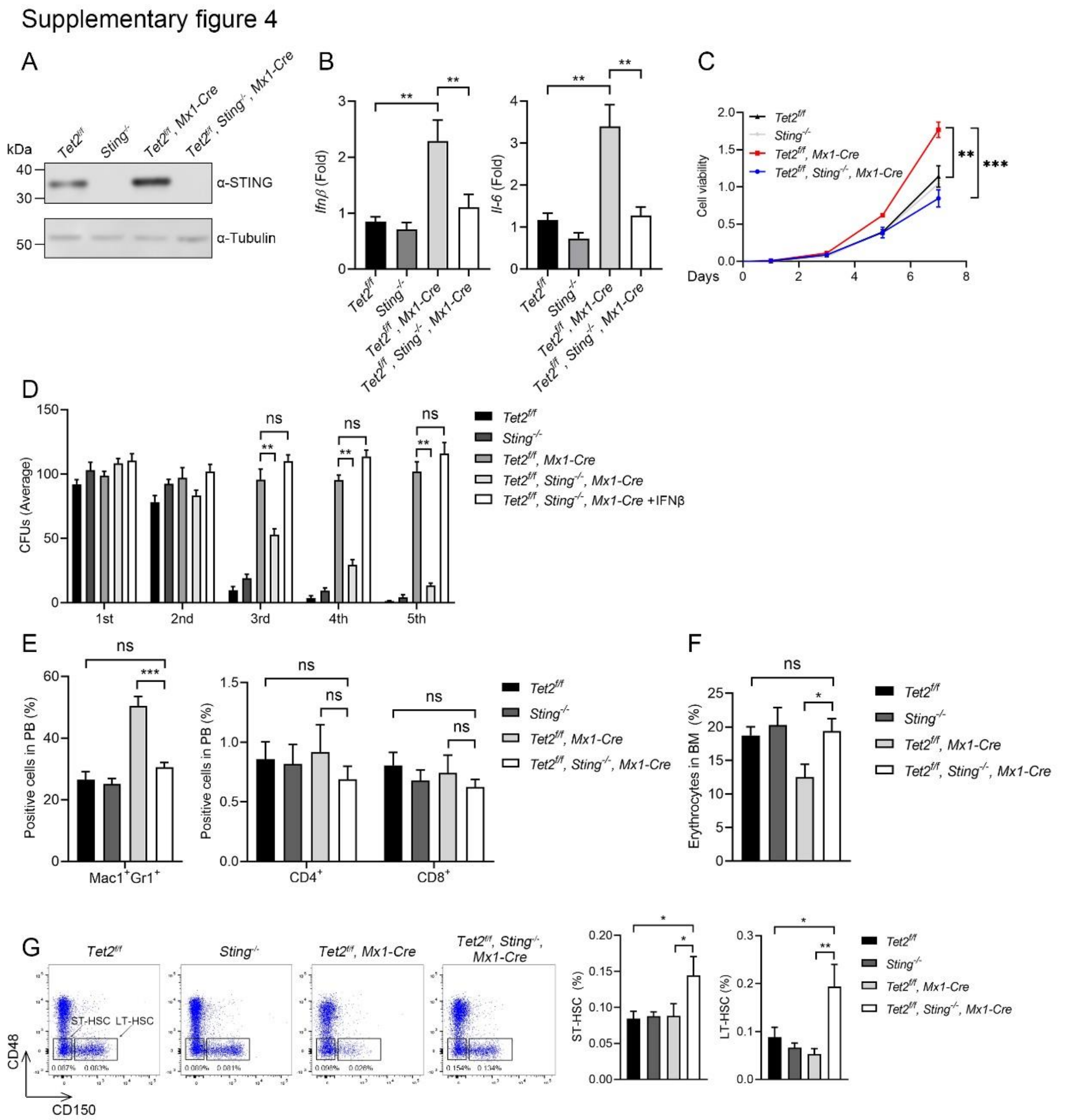


**Supplementary figure 4. *Sting* deletion attenuates increased growth advantage and myeloid-skewed differentiation induced by *Tet2* deficiency.**

(A) Western analysis of STING expression in the BM cKit$^+$ cells with indicated genotypes. Tubulin was used for normalization.

(B) *Sting* deletion mitigates the upregulation of *Ifnβ* and *Il-6* in *Tet2*$^{-/-}$ cKit$^+$ cells. The mRNA levels of *Ifnβ* and *Il-6* were quantified by qRT-PCR (n = 3 mice).

(C) *Sting* deletion reduces the growth rate of *Tet2*$^{-/-}$ hematopoietic progenitor cells. cKit+ cells were isolated from BM cells of 16-week-old mice and cultured in SFEM medium supplemented with SCF (100 ng/ml), IL-3 (10 ng/ml) and IL-6 (10 ng/ml). Cell viability was measured by CCK8 assay (n = 3 mice, biological replicates).

(D) Reduction in replating potential of isolated *Tet2*$^{-/-}$ LSK cells when lacking *Sting*.

IFNβ treatment (0.2 U/ml) restored the replating ability of *Tet2;Sting*$^{DKO}$ LSK cells (*n* = 3 experiments, each group with 1 mouse/experiment).

(E) *Sting* deletion attenuates *Tet2*-deficiency induced myeloid-skewed differentiation in the peripheral blood (PB). $CD4^+$ or $CD8^+$ T cells represent differentiation of lymphocytes (n = 5 mice).

(F) *Sting* deletion restores normal erythropoiesis of *Tet2*$^{-/-}$ mice (n = 5 mice). The proportion of $CD71^+Ter119^+$ erythrocytes was analyzed by FACS.

(G) The HSC pool was expended as a result of *Sting*, *Tet2* double deletion. Representative FACS images of HSCs are shown to the left. Quantification is shown to the right (n = 5 mice).

All mice (E-G) were analyzed between 40 to 44 weeks of age. Statistical significance was assessed by t-test (B-G). Data are mean ± s.e.m., *$P < 0.05$, **$P < 0.01$, ***$P < 0.005$, “ns”: not significant.

Supplementary figure 5

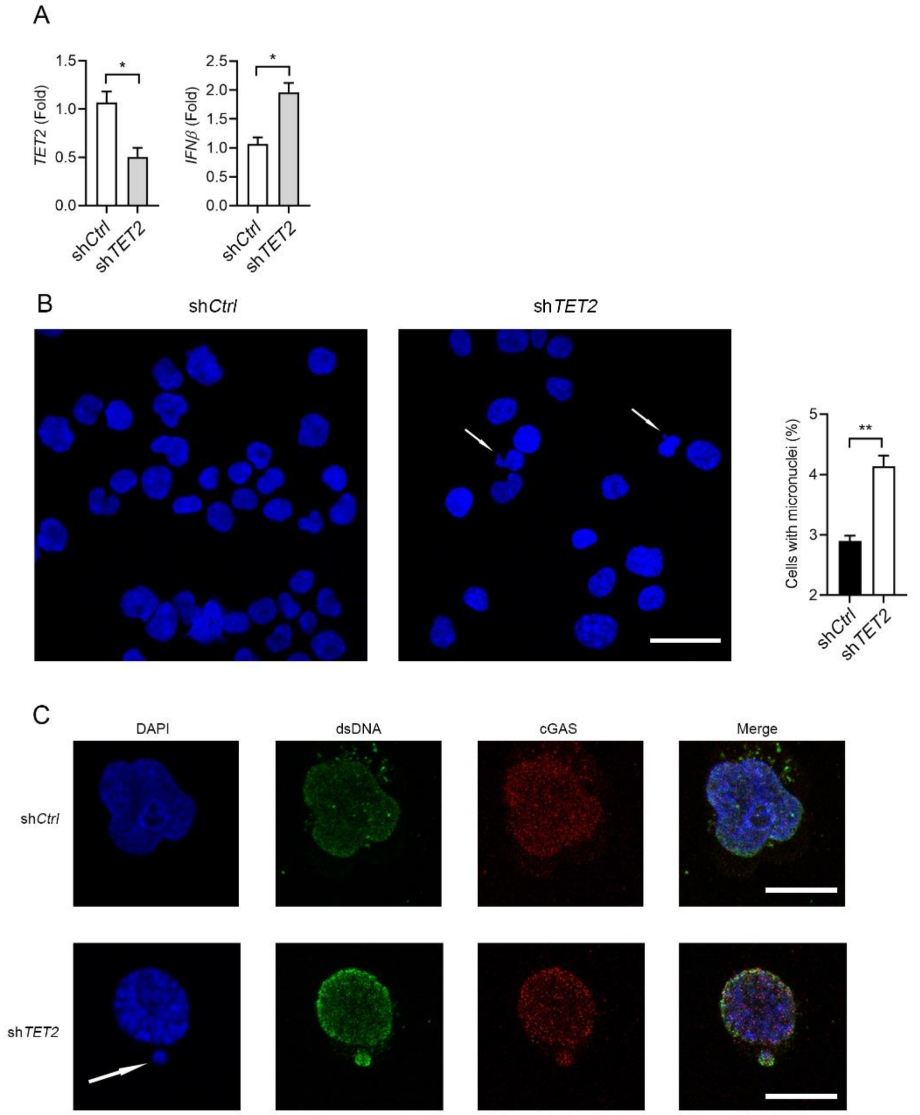


**Supplementary figure 5. cGAS-STING pathway is activated in TET2-ablated human HSCs**

(A) *TET2* and *IFNβ* expression in TET2-ablated CD34$^+$ cells quantified by qRT-PCR (n = 3 biological replicates). CD34$^+$ cells were isolated from cord blood and transduced with sh*TET2* or sh*Ctrl* lentivirus. The expression of *TET2* and *IFNβ* were analyzed at 72h after transducing.

(B) Accumulation of micronuclei in TET2-ablated CD34$^+$ cells revealed by DAPI staining. Arrows indicate micronuclei and the quantification is shown on the right (Scale bar, 50 μm).

(C) Immunofluorescence showing cGAS colocalizing with micronucleus and dsDNA in TET2-ablated $CD34^+$ cells. The arrow indicates micronucleus (Scale bar, 20 μm). cGAS (red) and dsDNA (green) signals were detected with specific antibodies. Statistical significance was assessed by t-test (A, B). Data are mean ± s.e.m. $*P < 0.05$; $**P < 0.01$; “ns”: not significant.

Supplementary figure 6

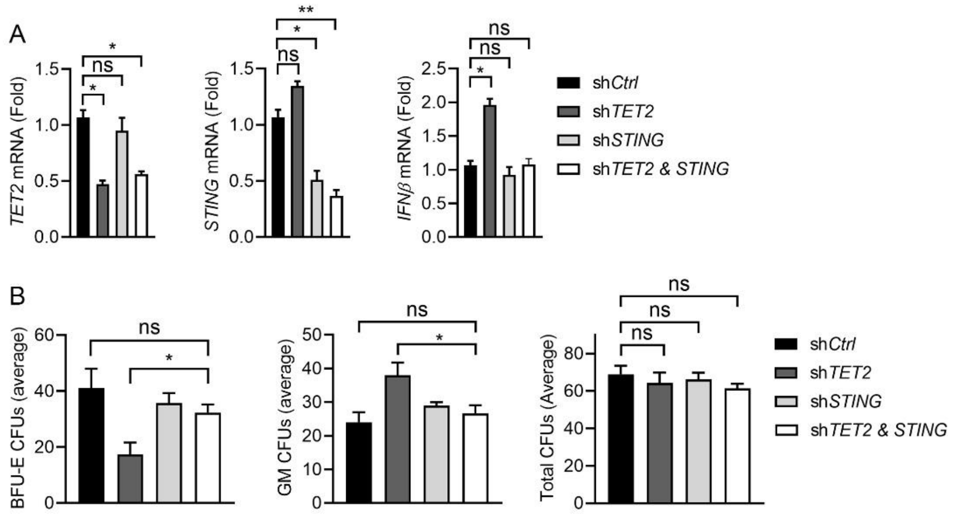


**Supplementary figure 6. Activated cGAS-STING pathway mediates skewed myeloid differentiation of TET2-ablated human HSCs in vitro.**

(A) STING depletion reduces the expression of *IFNβ* in TET2-ablated CD34$^+$ cells. The mRNA levels of *TET2*, *STING* and *IFNβ* were quantified by qRT-PCR (n = 3 biological replicates).

(B) STING depletion attenuates myeloid colony formation and restores erythroid colony formation of TET2-ablated CD34$^+$ cells (n = 3 biological replicates).

Statistical significance was assessed by t-test (A, B). Data are mean ± s.e.m. *$P <$ 0.05; “ns”: not significant.

Supplementary figure 7

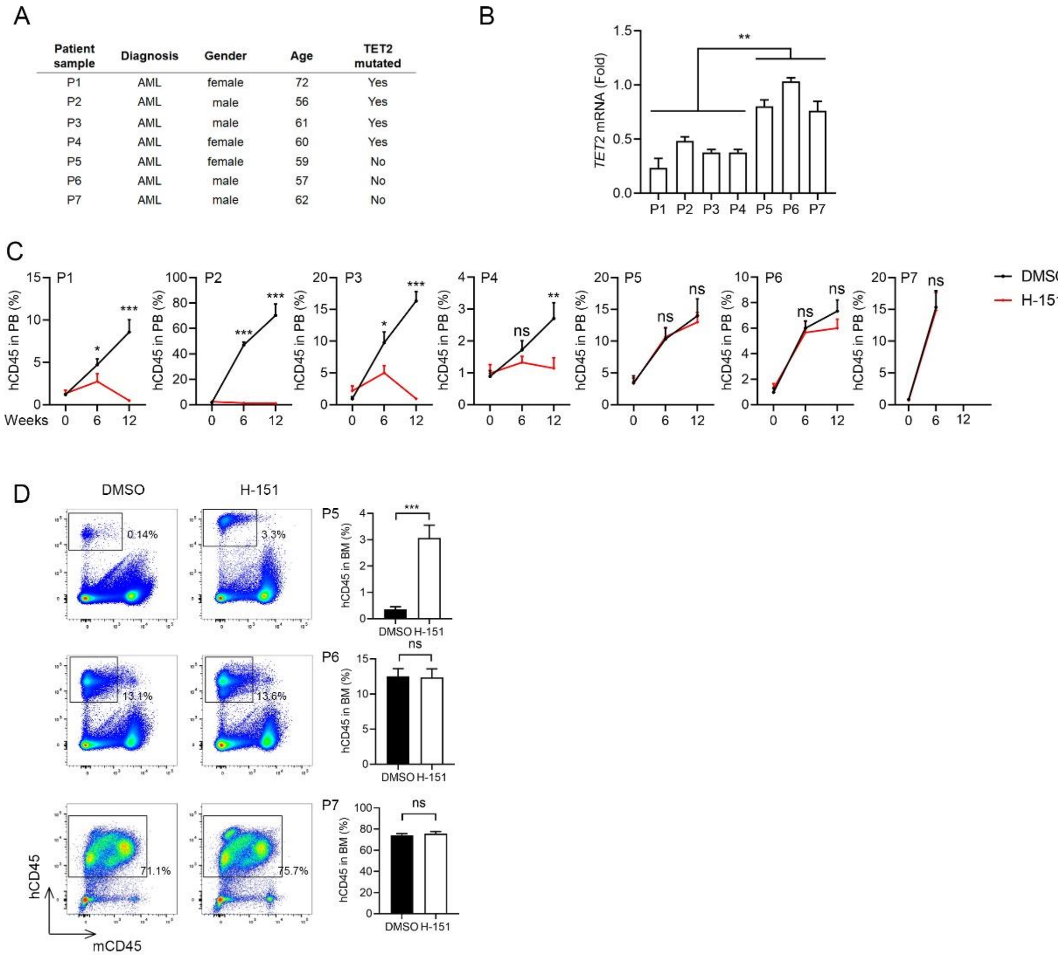


| Patient sample | Diagnosis | Gender | Age | TET2 mutated |
|---|---|---|---|---|
| P1 | AML | female | 72 | Yes |
| P2 | AML | male | 56 | Yes |
| P3 | AML | male | 61 | Yes |
| P4 | AML | female | 60 | Yes |
| P5 | AML | female | 59 | No |
| P6 | AML | male | 57 | No |
| P7 | AML | male | 62 | No |

**Supplementary figure 7. STING inhibition ameliorates the leukemogenesis of TET2-mutated mononuclear cells in AML patients-derived xenograft models.**

(A) AML patient information.

(B) *TET2* expression is downregulated in TET2-mutated AML patient cells. The mRNA levels of *TET2* were quantified by qRT-PCR (n = 3 biological replicates).

(C) STING inhibitor H-151 has no effect on the engraftment of TET2-normal patient AML cells in transplantation assay. Recipient-mice were treated with DMSO or H-151 through intraperitoneal injection every 3 days and the percentages of human CD45$^{+}$ cells in PB were analyzed at 14 weeks after transplantation (n = 3 mice).

(D) H-151 reduces the expansion of TET2-mutated patient AML cells in peripheral blood of B-NDG mice (n = 3 mice). Mice were treated with DMSO or H-151 through intraperitoneal injection every 3 days and the engraftments of human CD45$^{+}$ cells in peripheral blood were analyzed by FACS every 6 weeks. All three recipient mice of

P7 died at week 7.

Statistical significance was assessed with two-way ANOVA (B) and t-test (C, D). Data are mean ± s.e.m. $*P < 0.05$, $**P < 0.01$, $***P < 0.005$. “ns”: not significant.

### Supplementary methods

#### Lentivirus preparation

All shRNAs were expressed in pLKO.1-copGFP or pLKO.1-BFP lentivirus with the target sequences as follows, human *STING* (GCATCAAGGATCGGGTTTACA), human *TET2* (TATGAGTCTCGAACTCGCT), mouse *Sting* (GCATCAAGAATCGGGTTTATT), mouse *Mavs* (CCAGTGCTGATCTATTAGGAA), A pLKO.1 scramble control plasmid containing the following targeting sequence was used: CCTAAGGTTAAGTCGCCCTCG[1]. Lentiviral production was performed as previously described[2].

#### Colony formation assay

Lineage$^-$cKit$^+$Sca1$^+$ (LSK) cells were isolated from the BM cells of transgenic mice and plated into a 24-well plate with methylcellulose medium (M3434; StemCell Technologies) at 600 cells per well. Colony forming units were replated (1000 cells per well) every 7 days and colonies were scored at day 7 post-plating. To test the effect of C-178 and IFNβ, LSK cells were cultured in the presence of 0.5 μM C-178 and/or 0.2 unit/ml IFNβ. Human cord blood CD34$^+$ cells isolated through FACS were plated in methylcellulose medium (H4434, StemCell Technologies) at 600 cells per well and colonies were scored two weeks after plating. Mononuclear cell (MNC) fractions were obtained from BM of patients with AML by density gradient centrifugation using Ficoll-Paque Plus (GE Healthcare). MNCs were plated into methylcellulose (H4434, StemCell Technologies) at $1 \times 10^4$ cells per well in the presence of DMSO or 1 μM H-151. Plates were incubated at 37 °C at 5% $CO_2$, and colonies were counted two weeks after plating.

#### Transduction of human cord blood CD34$^+$ cells and mouse cKit$^+$ cells

Human cord blood CD34$^+$ cells were enriched using a magnetic bead sorting system according to the manufacturer’s instructions (Miltenyi Biotec, Bergisch Gladbach, Germany). For the initial expansion, cord blood CD34$^+$ cells were cultivated in StemSpan SFEM (09650, StemCell Technologies) supplemented with 100 ng/ml human stem cell factor (SCF), 50 ng/ml Fms-like tyrosine kinase 3 ligand (Flt-3L) and 50 ng/ml thrombopoietin (TPO) for 1 day. The pre-stimulated $CD34^+$ cells were infected with lentivirus carrying shRNA targeting *TET2*, *STING* or a control vector in the presence of polybrene (5 µg/ml, Sigma). The positive cells were sorted on the flow cytometer (FACS Aria III, BD Biosciences, San Jose, CA) 72 hours post-infection. For colony-forming assays, the sorted $GFP^+BFP^+$ cells were cultured in methylcellulose (H4434, StemCell Technologies) supplemented with 10% IMDM (Sigma) basic culture medium for 14 days.

Mouse BM LSK cells harvested from transgenic mice were cultured overnight in StemSpan SFEM supplemented with mouse recombinant SCF (100 ng/ml), IL-6 (10 ng/ml) and IL-3 (10ng/ml). The next day, cells were infected with lentivirus carrying shRNA targeting *Sting, Mavs* or an empty vector control in the presence of polybrene (5 µg/ml, Sigma) and centrifuged at 1000g, 32℃ for 1 hour. The spin infection was repeated the next day. Forty-eight hours after transduction, the GFP positive cells were sorted, and $1 \times 10^5$ cells were lysed to determine the knockdown efficiency and cytokine expression.

**Flow cytometric analysis and cell sorting**

Single-cell suspensions from BM and PB were stained with fluorochrome-conjugated antibodies. For lineage stratification, cells from PB and BM were stained with FITC-Gr1, PE-cy7-Mac1, FITC-CD4, PE-cy7-CD8, PE-Ter119 and APC-CD71 for 30min at 4°C. For LT-HSCs and ST-HSCs, BM cells were stained with FITC-conjugated lineage cocktail (B220, CD4, CD5, CD8, Gr1 and Ter119), Percp-cy5.5-Sca1, BV421-cKit, PE-CD150, APC-CD48. LT-HSC were immunophenotypically defined as $Lin^-cKit^+Sca1^+CD48^-CD150^+$ and ST-HSCs as $Lin^-cKit^+Sca1^+CD48^-CD150^-$. For committed progenitor cells, granulocyte-monocyte progenitors (GMP) were immunophenotypically defined as $Lin^-cKit^+$ $Sca1^-CD34^+CD16/32^{high}$, common myeloid progenitors (CMP) as $Lin^-cKit^+Sca1^-CD34^{int/+}CD16/32^{int}$, megakaryocyte-erythrocyte progenitors (MEP) as $Lin^-cKit^+Sca1^-CD34^-CD16/32^{low}$) and common lymphoid progenitors (CLP) as $lin^-cKit^+Sca1^{int}$ $CD127^+$. The analyses and sorting were performed using a BD FACScanto II cytometer or a BD FACSAria™ III cell sorter. All data were analyzed using FlowJo software, version 10.

**Immunofluorescence assay**

FACS-sorted mice hematopoietic cells and human cord blood $CD34^+$ cells were concentrated using a cytospins for immunofluorescence analysis. Immunofluorescence was performed as previously described[5]. The following primary antibodies were applied: anti-phospho-H2AX (Ser139) (Millipore, 05-636), anti-ds DNA (Abcam, ab27156), anti-cGAS (CST 79978). Images were acquired using a Leica TCS SP8 confocal laser microscopy system.

**PCR and Real-time quantitative PCR**

PCR Genotyping primers were as follows: *Tet2*: 5'-ACTCATTAGTGAAATATGTGAGTG-3', 5'-CTGCTTAGTTCAATGCCAACC-3', 5'-ACACAGAGAAAAGGGTACGTGAA-3'. Wild-type band = 447 bp, floxed band = 536 bp, knockout band = 617 bp. *Sting*: 5'-TGCTGTAGGATGCTATGTGC-3', 5'-ACAGAGGGTTACCTGGACTG-3'. Wild-type band = 224 bp, floxed band = 312 bp, knockout band = 227 bp.

Total RNA was extracted with TRIzol Reagent (Invitrogen, Thermo Fisher Scientific), and cDNA was synthesized using HyperScript III 1st Strand cDNA Synthesis Kit with gDNA Remover (EnzyArtisan, R201) according to the manufacturer's instructions. Real-time quantitative PCR was performed with S6 Universal SYBR qPCR Mix (EnzyArtisan, Q204) on a LightCycler 480 II (Roche Applied Science). Expression of genes of interest were normalized to the housekeeping gene Actb using the 2–ΔΔCt method. qPCR primers were as follows, mouse *Ifnβ*: 5'-CCCTATGGAGATGACGGAGA-3', 5'-CTGTCTGCTGGTGGAGTTCA-3'. Mouse *Il-6*: 5'-TCCATCCAGTTGCCTTCTTG-3', 5'-GGTCTGTTGGGAGTGGTATC-3'. Mouse *Actin*: 5'-TGACGTTGACATCCGTAAAGACC-3', 5'-AAGGGTGTAAAACGCAGCTCA-3'. Mouse *Sting*: 5'-GTCTAGGAAGCAGAAGATGCCA-3', 5'- GAGGACCAGAAGGCCAAACA-3'. Mouse *Mavs*: 5'-CTGCCTCACAGCTAGTGACC-3', CCGGCGCTGGAGATTATTG. Human *TET2*: 5'-GGAAAGCTTTTCAGCTGCAGC-3', 5'-CTTGCACAACATGCAGAATGGCAGC-3'. Human *GAPDH*: 5'-AGAAGGCTGGGGCTCATTTG-3', 5'-AGGGGCCATCCACAGTCTTC-3'. Human *IFNβ*: 5'- CAGCAGTTCCAGAAGGAGGA-3', 5'-

AGCCAGGAGGTTCTCAACAA-3’. Human STING: 5’-CCTTGGTTCTGCTGAGTGCC-3’, 5’-CCGGTACCTGGAGTGGATGT-3’.

**RNA-seq library preparation**

About 300 hematopoietic cells per group were sorted directly into Smart-seq2 lysis buffer by FACS. Sorted cells were lysed and reverse-transcribed RNA was amplified to obtain enough cDNA by a modified SMART-Seq2 protocol[3,4]. cDNA was quantified by Qubit 3 (Invitrogen, Q33327), and 5 ng cDNA was used for cDNA library construction with TruePrep DNA Library Prep Kit V2 for Illumina (Vazyme, Cat. TD502).

**RNA-seq data analysis**

The raw pair-end RNA-seq FASTQ data were trimmed to remove low-quality bases and adaptor sequences by Trim Galore (v0.5.0) with default settings. Then, the clean RNA-seq FASTQ data were mapped to mouse reference genome mm10 using Hisat2 (v2.1.0) with default parameters. Differentially expressed gene (DEG) analysis was performed by using DESeq2 R package with the raw count. Only genes with adjusted P value less than 0.05 and at least 2-fold-change were considered to be differentially expressed. DEGs were submitted to DAVID 6.7 (https://david.ncifcrf.gov) for gene ontology (GO) and KEGG pathway enrichment analyses.

**Western Blot Analysis**

Equal numbers of cells from each population were isolated with FACS or LS column (Miltenyi Biotec. Cat. No. 130-042-401). Cells were collected and resuspended in lysis buffer containing Tris 8.0 20mM, NaCl 150mM, Triton X-100 0.5%, protease inhibitor cocktail (Roche). BM-MNCs were harvested from mice with different genotypes. Lysates of BM-MNCs were prepared with lysis buffer (Tris-HCl 7.5 20mM, NaCl 150mM, Triton X-100 0.5%, SDS 0.2%). Lysates were subjected to ultrasonication for 5 min to disrupt the genomic DNA and then mixed with SDS loading buffer. Blots were developed with ECL detection reagent (180-5001, Tanon) and imaged on a CCD imager (GE Healthcare).

**Reagents and antibodies**

STING inhibitor C-176, C-178, H-151 were purchased from MCE (Cat. No. HY-112906; HY-123963; HY-112693). 2’3’-cGAMP (Cat. No. SML1299) and ATP-$^{13}C_{10}$,$^{15}N_5$ (Cat. No. 645702) were obtained from Sigma. All other chemical reagents were purchased from Sigma or Sangon Biotech. Flag antibody(F3165), Tubulin antibody (SAB4500088), goat anti-rabbit (AP132) and goat-anti mouse secondary antibodies (AP124) were purchased from Sigma; all FACS antibodies were purchased from Invitrogen or BD; antibodies against MAVS (83000), STING (13647), p-STING (62912), p-TBK1 (5483) were obtained from CST.

**Generation of $^{13}C_{10}{}^{15}N_5$-labeled cGAMP**

Recombinant His-SUMO-cGAS (human) was expressed and purified in *E. coli* strain Rosetta (DE3) as previously described[5]. For $^{13}C_{10}{}^{15}N_5$-labeled cGAMP in vitro synthesis, 100 μg of His-SUMO-cGAS protein was mixed with Buffer R (20 mM HEPES, pH 7.2, 5 mM $MgCl_2$, 1 mM ATP-$^{13}C_{10}$,$^{15}N_5$, 1 mM GTP, and 0.1 mM EGTA) in the presence of 0.1 mg/ml HT-DNA. The mixture was incubated at 37°C for 1 hour, then heated at 95°C for 5 min, centrifuged at 10000g, 4°C, for 10 min. The $^{13}C_{10}{}^{15}N_5$-labeled cGAMP was in the heat-resistant supernatant and was used as an internal standard for the quantification of cellular cGAMP.

**Extraction of endogenous cGAMP in BM-MNCs**

Total BM-MNCs (1 mouse) were collected and washed twice with cold PBS, resuspended in cold 80% (vol/vol) methanol with 2% (vol/vol) acetic acid (HAc), and stored at −80 °C. On the day of analysis, 50 fmol $^{13}C_{10}{}^{15}N_5$-labeled cGAMP (+15 atomic mass units) was added into the BM cells as internal standard. Cells were homogenized for 20 s and centrifugated at 12000g for 10 min. The pellets were extracted in 20% (vol/vol) methanol and 2% HAc, and all supernatants were pooled. The supernatants were then loaded onto HyperSep Aminopropyl SPE Columns (Thermo Scientific) to remove impurities and enrich cGAMP. Briefly, the columns were washed sequentially with 1 ml methanol and 500 μl 2% HAc for two times; then, all supernatants were loaded onto SPE columns (about 1ml); after drawing through the extracts, columns were washed twice with 500μl 2% HAc and once with 500μl 80% methanol; cGAMP enrichments were eluted with 4% (vol/vol) ammonium hydroxide in 80% methanol. The eluents were spin-vacuumed to dry and reconstituted in 40% acetonitrile: 40% methanol: 20% $H_2O$ (vol/vol). The eluents were cleared by centrifugation and transferred to autosampler vials for MS analyses.

**Quantification of cGAMP with LC-MS**

The LC-MS/MS analysis was performed on an Agilent 1290 Infinity II LC System equipped with a ACQUITY UPLC BEH Amide column (1.7 µm, 2.1 × 150 mm; Waters) coupled by ESI to a QTRAP 6500$^+$ (AB Sciex). The column was maintained at 35°C and eluents were injected at a constant flow rate of 300 µl/min. The binary mobile phase was composed of 0.125% formic acid in 50/50 (vol/vol) acetonitrile/water (A) and 90/10 (vol/vol) acetonitrile/water (B), and samples were flowed through the following gradient: 0 min, 99% B; 3 min, 99% B; 5 min, 50% B; 5.5 min, 35% B; 10.5 min, 30% B; 11 min, 5% B; 14.5min, 5% B; 15 min, 99% B, and 20 min, 99% B. The mass spectrometer was operated in the positive ion mode with the following settings: ion spray voltage: +5500 V, declustering potential: 85 V, entrance potential: 10 V, collision energy: 32 V, source temperature: 550°C, and curtain, ion source gas at 35, 55. cGAMP and the $^{13}C_{10}{}^{15}N_5$-labeled cGAMP were detected in multiple reaction monitoring (MRM) mode for four mass transitions, respectively (cGAMP: 675-136, 675-152, 675-476, and 675-524; and the $^{13}C_{10}{}^{15}N_5$-labeled cGAMP: 691-146, 691-152, 691-491, and 691-539). Peak identities were verified by the analysis of commercial cGAMP and $^{13}C_{10}{}^{15}N_5$-labeled cGAMP spike-in and selected areas were integrated with the Sciex OS software 2.0. Relative abundance of endogenous cGAMP was calculated based on the difference of peak aeras between the control and *Tet2*-deficient groups using a calibration curve, and was normalized to internal standard.

**Comet assay**

Mouse BM LSK cells were collected and resuspended in phosphate-buffered saline (PBS) at $1 \times 10^5$ cells per ml. The comet assay was performed according to the manufacturer's instructions (4250-050-K; Trevigen, *Gaithersburg*, Maryland). DNA damage was measured by tail moments using comet-score software.

**Statistical analysis**

Data are shown as mean ± s.e.m. To assess the statistical significance, we used unpaired Student's t-tests for comparison between two groups and two-way (more than two genotypes) ANOVA for multiple groups. Survival curves were compared using Mantel-Cox log-rank test. All statistical tests were performed using GraphPad Prism software.
